# Relative phase of membrane potential theta oscillations between individual hippocampal neurons code space

**DOI:** 10.1101/2025.11.14.688496

**Authors:** Mohamed Athif, Samantha Malmberg, Rebecca Mount, Kush Patel, Yangyang Wang, Dana Shaw, Cara Ravasio, Sheng Xiao, Enrico Rillosi, Hua-an Tseng, Sarah E. Plutkis, Jonathan B. Grimm, Luke D. Lavis, Jerome Mertz, Xue Han, Michael E. Hasselmo

**Affiliations:** Department of Biomedical Engineering, Boston University, Boston, MA; Department of Psychological and Brain Sciences, Boston University, Boston, MA; Center for Systems Neuroscience, Boston University, Boston, MA; Graduate Program in Neuroscience, Boston University, Boston, MA; Undergraduate Program in Neuroscience, Boston University, Boston, MA; Janelia Research Campus, Howard Hughes Medical Institute, Ashburn, VA

**Keywords:** Voltage imaging, membrane potential, subthreshold dynamics, Voltron2, phase procession

## Abstract

The timing of spikes dictates a neuron’s impact on downstream circuits and behavior, and spike timing is determined by the membrane potential (Vm). However, due to technical challenges, it has been impossible to analyze the relative timing of Vm dynamics between neurons during behavior. Using large scale membrane voltage imaging, we simultaneously recorded Vm from many individual hippocampal neurons in animals engaged in a virtual spatial task. We found that relative phase of Vm theta oscillations across neurons exhibit gradual or discrete shifts depending on spatial position. This finding extends beyond previous studies showing Vm dynamics in single neurons or spiking activity in multiple neurons, revealing previously unknown evidence for consistent coding of space by spike-independent relative phase of Vm theta dynamics between neurons.

## INTRODUCTION

The dynamics of neuronal membrane potential (Vm) underlie the processing of information by individual neurons. Vm integrates network influences from synaptic conductances in conjunction with intrinsic voltage-dependent conductances to determine whether a neuron will fire an action potential and influence the subsequent behavior of the animal (Koch & Segev, 2000; Izhikevich, 2006). However, most studies of neuronal activity in behaving animals have focused on the timing of spiking activity rather than Vm dynamics (O’Keefe & Recce, 1993; Wilson & McNaughton, 1993; Skaggs et al., 1996; Huxter et al., 2003; Vollan et al., 2025). The focus on spiking is partly due to the challenges of recording Vm using intracellular recording with micropipettes in behaving animals, which only allow for recording of single neurons for limited durations (Harvey et al., 2009; Epsztein et al., 2011; Domnisoru et al., 2013; Schmidt-Hieber & Häusser, 2013; Bittner et al., 2015, 2017; Grienberger et al., 2017).

What is known about Vm in limbic structures of intact animals comes from intracellular recordings of single neurons, which show subthreshold Vm oscillations of membrane potential at theta frequency in single neurons of the hippocampus (Fujita & Sato, 1964; Leung & Chi Yiu Yim, 1986; Kamondi et al., 1998; Harvey et al., 2009; Epsztein et al., 2011) and entorhinal cortex (Domnisoru et al., 2013; Schmidt-Hieber & Häusser, 2013). Intracellular recording of single neurons also shows that shifts in theta phase of Vm relative to network local field potential (LFP) oscillations can code location (Harvey et al., 2009; Domnisoru et al., 2013). However, because of technical challenges, no-one has analyzed the relative Vm phase of multiple simultaneously recorded neurons relative to each other rather than the LFP. The LFP reflects a mean phase across the network, so using LFP as a phase reference could smear out phase information of individual neurons and prevent detection of coding based on relative phase of Vm across pairs of individual neurons. Recent innovations in genetically encoded voltage indicators permit simultaneous imaging of Vm dynamics in multiple neurons (Piatkevich et al., 2019; Lowet et al., 2021, 2022, 2023; Xiao et al., 2021, 2024; Tseng et al., 2022; Abdelfattah et al., 2023; Shroff et al., 2023) enabling the analysis of continuous relationship of Vm between neurons during behavior.

While previous studies showed the importance of spike to LFP phase coding in the hippocampus, spiking provides a discretized sampling of membrane potential effects. The spiking of hippocampal neurons shows strong relationships to the phase of LFP theta oscillations (Buzsáki et al., 1983; Fox et al., 1986; O’Keefe & Recce, 1993; Skaggs et al., 1996; Huxter et al., 2003) and many neurons code location by theta phase precession, in which the spikes shift to earlier phases of LFP theta oscillations as animals navigate through the firing field of a hippocampal place cell (O’Keefe & Recce, 1993; Skaggs et al., 1996; Huxter et al., 2003) or an entorhinal grid cell (Hafting et al., 2008; Climer et al., 2013). Importantly, spiking phase coding for location can be independent of firing rate, as shown in lateral septum cells that fire throughout the environment but at different phases for different locations (Tingley & Buzsáki, 2018). But these studies do not reveal the potential coding mechanisms of subthreshold Vm.

To probe spike-independent Vm phase coding of behavior, we performed large scale voltage imaging from multiple individual CA1 neurons simultaneously using the genetically encoded voltage sensor Voltron2 (Abdelfattah et al., 2023) and a targeted illumination microscope (Piatkevich et al., 2019; Lowet et al., 2021, 2023; Xiao et al., 2021, 2024; Tseng et al., 2022), while head-fixed mice were performing a virtual spatial task. We analyzed how the relative phase of Vm between different CA1 neurons consistently shifts based on positions on the virtual track. We show that coding by relative phase of Vm oscillations between neurons can occur even on cycles when the neurons are not spiking, and coding by relative phase can occur in neurons without localized firing fields. These results reveal previously unknown evidence for consistent coding of position by the relative phase of Vm theta oscillations in different neurons.

## RESULTS

### Large scale voltage imaging of CA1 neurons during a virtual spatial navigation task

To probe Vm phase coding in the hippocampus, we performed large scale voltage imaging from many CA1 neurons simultaneously in head-fixed mice unidirectionally navigating in a looped virtual reality (VR) track. The task is similar to that used in previous studies, which showed spiking of place fields that are distributed over the full track quantified with electrophysiology (Chen et al., 2013) or calcium imaging (Dombeck et al., 2010; Robinson et al., 2020). Each 200 cm VR loop contains various visual cues and a 25 cm reward zone marked with distinct floor patterns (Figure 1A). To perform cellular voltage imaging, we surgically implanted an imaging window over CA1 and virally transduced CA1 neurons with the voltage sensor Voltron2 (Abdelfattah et al., 2023). During each imaging day, mice performed the VR task for 15 minutes, while Vm of multiple CA1 neurons was recorded with a high-speed custom targeted illumination microscope (Xiao et al., 2021) (Figure 1B). Over days, mice progressively completed more traversals (trials) per day, and consistently improved their behavioral performance quantified as lick selectivity that measures the preference of licking within the reward zone versus the opposite zone defined as the 25 cm non-rewarded middle part of the loop (Figure 1C, D).

During each recording day, Voltron2 fluorescence was recorded at ∼800 frames per second. We then motion corrected each image stack offline, manually segmented individual neurons, extracted Vm, and identified spikes (Methods, Figure 2A). We routinely imaged from 13 ± 9 neurons simultaneously (mean ± standard deviation, n=2076 neurons from 174 sessions in 7 mice, Figure 2A and Supplementary table 1). Vm and spiking were further manually inspected to confirm minimal motion, and sufficient spike-to-baseline ratio (Figure 2B, C, and Supplementary figure 1).

**Figure 1:**
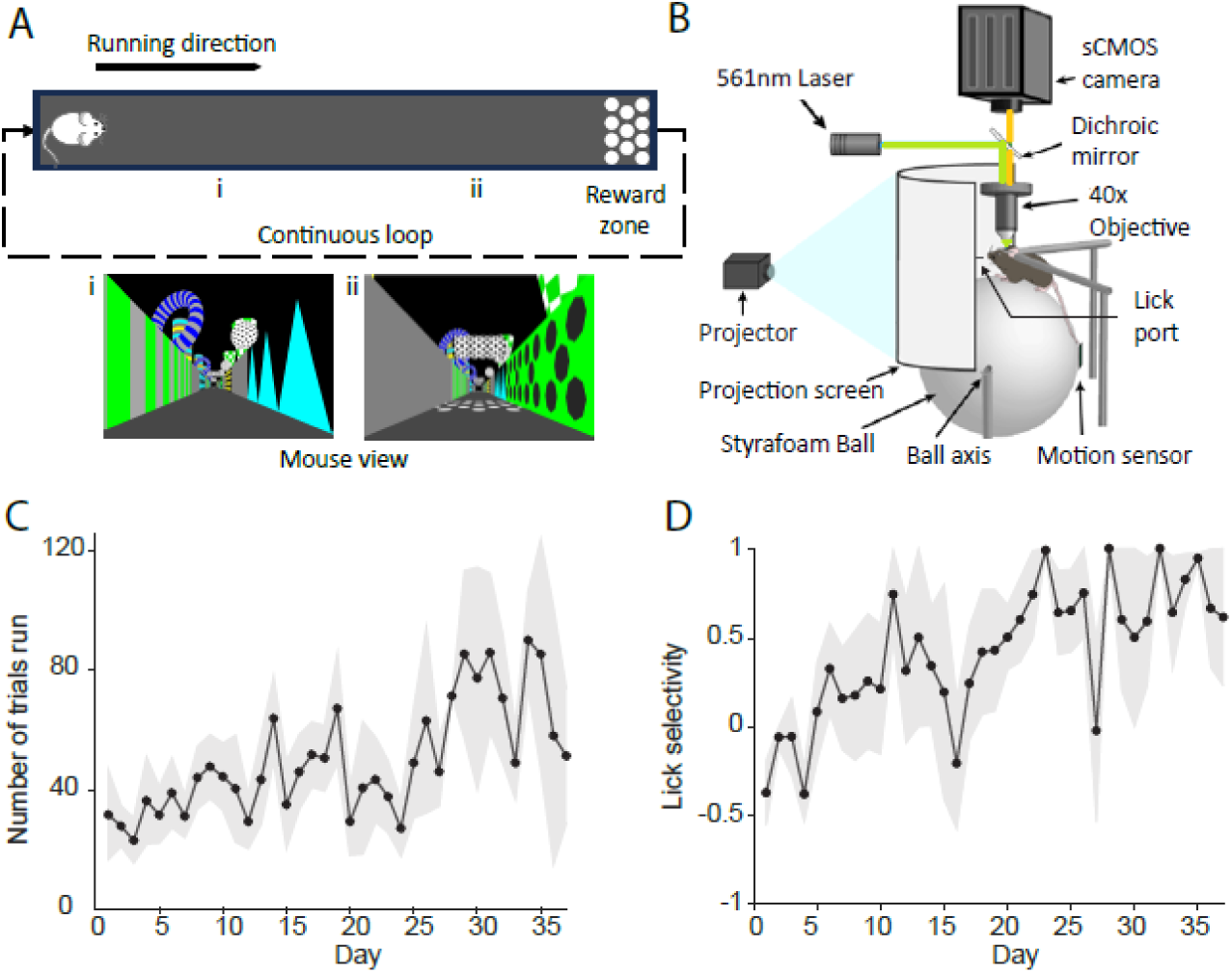
Voltage imaging of CA1 neurons in mice performing a virtual reality (VR) spatial task. (A) Top, illustration of the behavioral task. A mouse running unidirectionally in a continuously looped VR world with proximal cues (wall patterns) and distal cues (cylinder/ sphere/ donut), and a reward zone marked by unique floor patterns. Bottom, two snapshots of the VR environment (i) further away or (ii) close to the reward zone. (B) Illustration of the experimental setup, showing a head-fixed mouse on a spherical treadmill facing a screen displaying the VR world under a microscope for voltage imaging. (C) The number of trials mice completed during each recording day throughout training (mean ± S.E.M, n = 7 mice) (D) Lick selectivity across training days (mean ± S.E.M, n = 7 mice). Lick selectivity = (reward zone licks - opposite zone licks) / (reward zone licks + opposite zone licks).

**Figure 2.**
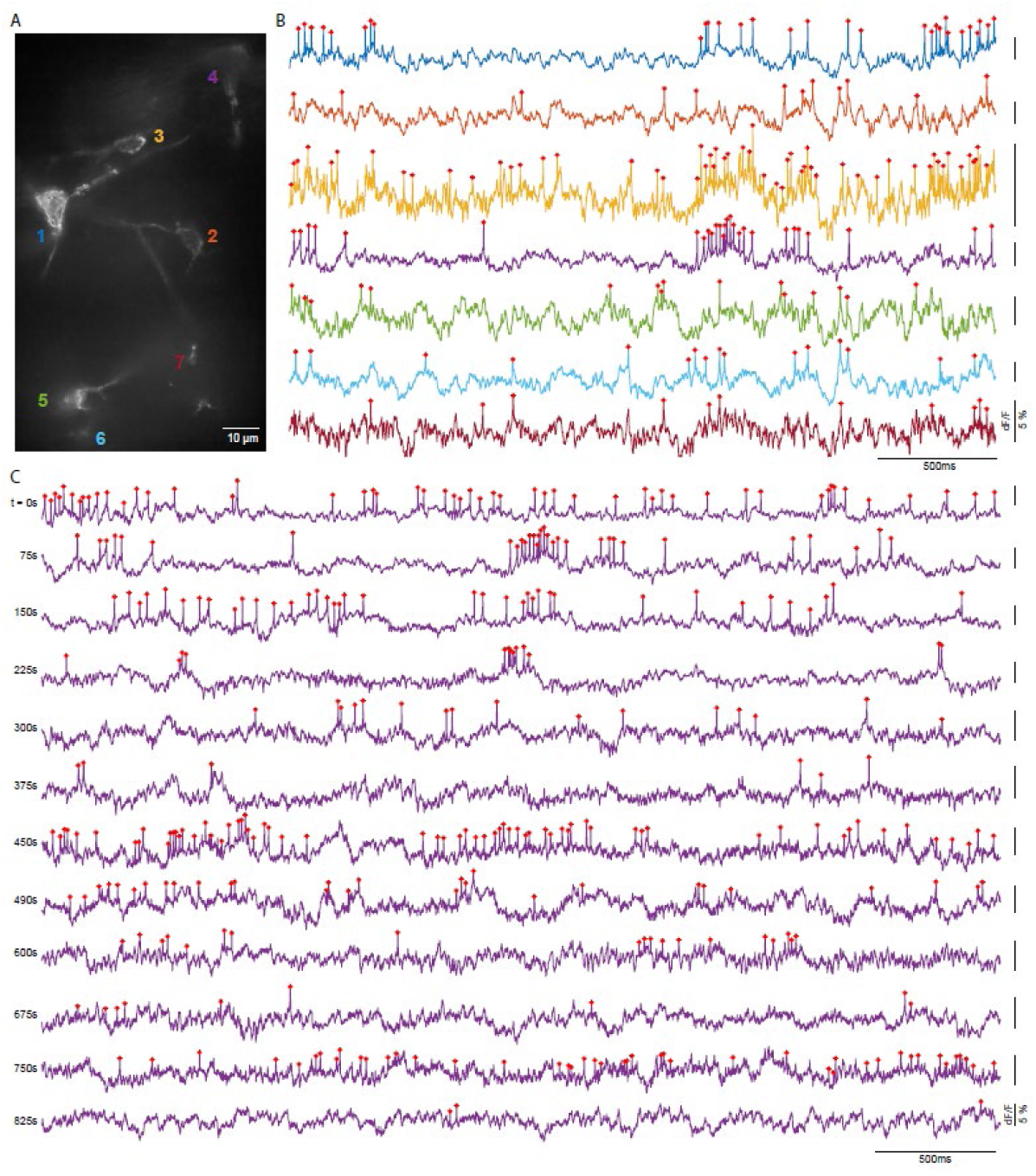
Simultaneous voltage imaging of multiple CA1 neurons while mice were performing the VR spatial task. (A) An example imaging field of view, showing 7 simultaneously recorded neurons. (B) Example Vm fluorescence traces from all 7 cells shown in A, at 75s from the start of the experiment. The color of the trace corresponds to the color of the neuron number shown in A. (C) Example Vm traces of neuron 4 throughout a 15-minute-long recording session. Each row shows an example 4 second Vm from the continuous recording. Red asterisks denote identified spikes.

### Vm theta oscillations in simultaneously recorded CA1 neurons exhibit prominent and stable phase differences during spatial navigation

CA1 neurons are known to exhibit prominent Vm theta rhythmicity, and Vm power spectrum analysis confirmed that most neurons had dominant theta oscillations (Figure 3B,D). We then detected spikes from Vm traces, and observed that spiking in most neurons predominantly occurred on the rising phase of Vm and LFP theta oscillations, consistent with previous intracellular electrophysiological (Noguchi et al., 2023) and voltage imaging (Kannan et al., 2022; Lowet et al., 2023) studies (Supplementary figure 2). Furthermore, we identified place cells with spatially specific spiking that tiled the linear track (Supplementary figure 3), and confirmed that some place cells exhibit the characteristic phase precession of spike times relative to LFP theta oscillations as broadly reported previously (Supplementary figure 4). Finally, we evaluated the temporal relationship between Vm and LFP theta oscillations (6-10 Hz), and confirmed that Vm theta peak in many cells also occurred at specific phases of LFP theta, though the exact temporal relationship between Vm and LFP theta varied across neurons (Supplementary figure 5).

In addition to capturing spiking, voltage imaging can record intracellular Vm from multiple neurons simultaneously. Thus, we further explored the temporal relationship of Vm theta oscillations between simultaneously recorded neuron pairs. We first identified the peak of each theta cycle in individual neurons by filtering Vm at 6-10 Hz. Aligning the Vm traces of one neuron around its own theta peaks confirmed that most neurons exhibit prominent Vm theta time scale fluctuations, with a 2^nd^ peak occurring at ∼160ms from the peak (Figure 3A, C, Ei). Power analysis of the full recorded traces (Figure 3B, 3D) and mean aligned Vm confirmed the theta rhythmicity in most neurons (Figure 3Fi) .

Interestingly, when aligning to the theta peaks of another simultaneously recorded neuron, we noted large heterogeneity of the corresponding peak-aligned Vm. While theta oscillations in most neuron pairs peaked around the same time with small temporal shifts of a few milliseconds (Figure 3Eii, iv, v), some pairs were almost completely antiphase with opposite peaks (Figure 3Eiii). Across the 41,553 pairs of neurons recorded, most had Vm peaks around the same time (centered at 0 or 360 degrees), Figure 3Fii), but a significant proportion had peaks distributed across a wide phase range with many exhibiting almost antiphase (antiphase is around 180 degrees, Figure 3Fii).

**Figure 3:**
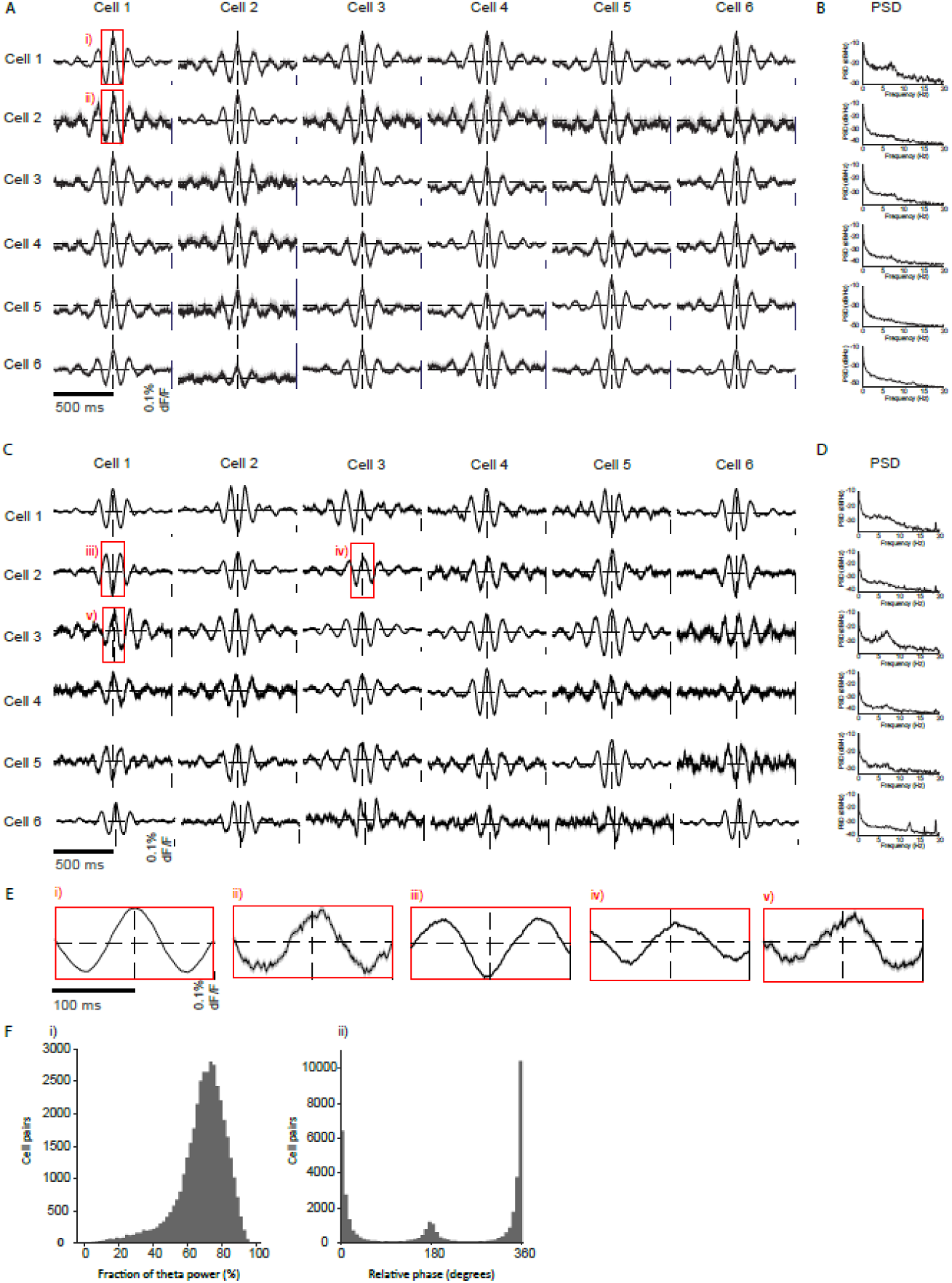
Relative Vm theta phase differences between simultaneously recorded neuron pairs. (A) Vm of a neuron aligned to the peak of its own Vm theta oscillations or the peak of theta oscillation in other simultaneously recorded neurons. This recording session had 6 neurons. Shown are mean ± S.E.M. Dashed lines indicate 0. (B) Vm power spectral density of the corresponding neuron shown in A. (C, D) Same as A and B, but for another recording session with 6 neurons. (E) Zoom-ins of highlighted boxes in A (i, ii) and B (iii, iv, v). (Fi) The distribution of theta power of the mean Vm aligned to theta peaks across all neurons analyzed (n=2,076 neurons). (Fii) Histogram of the relative difference of Vm theta peaks across all neuron pairs. (n=41,553 pairs).

### Relative Vm theta phase between neurons codes spatial position

As spiking activity in hippocampal neurons is known to precess relative to LFP theta oscillations, we next examined whether the relative Vm theta phase between CA1 neuron pairs encodes space. We first identified the time of each theta peak of one neuron within a pair, and then located the corresponding spatial location of the mouse and computed the relative phase shift of the pair as the instantaneous relative theta phase of the other neuron. To visualize the relationship of phase shift with spatial location, we plotted 2-dimensional histograms of relative theta phase aligned to the start of the linear track (Fig. 4). The stability of these 2D histograms was confirmed by comparing the theta cycles within the first half versus the last half of the recordings. This led to 34,560 2D histograms that exhibited stable phase relationships across all recording sessions in all mice. These 2D histograms were then clustered into 5 distinct groups using K-Means Clustering algorithm (Methods).

The major cluster, cluster A, contained 35.8% of the pairs. In this cluster, relative phase of neuron pairs dynamically changed along the spatial position, suggesting that the relative phase between neuron pairs codes space (Figure 4A, Supplementary figure 6). In clusters B (15.5% of pairs) and D (27.7% of pairs), the relative phase was centered at 180 degrees throughout the spatial location, with cluster B exhibiting a tighter phase angle than cluster D (Figure 4B, D, F). In cluster C (15.4% of pairs), the relative phase is tightly centered at 0 degree (Figure 4C). The smallest cluster (5.5% of pairs), Cluster E, contains pairs that do not have strong phase-position relationships (Figure 4E).

**Figure 4:**
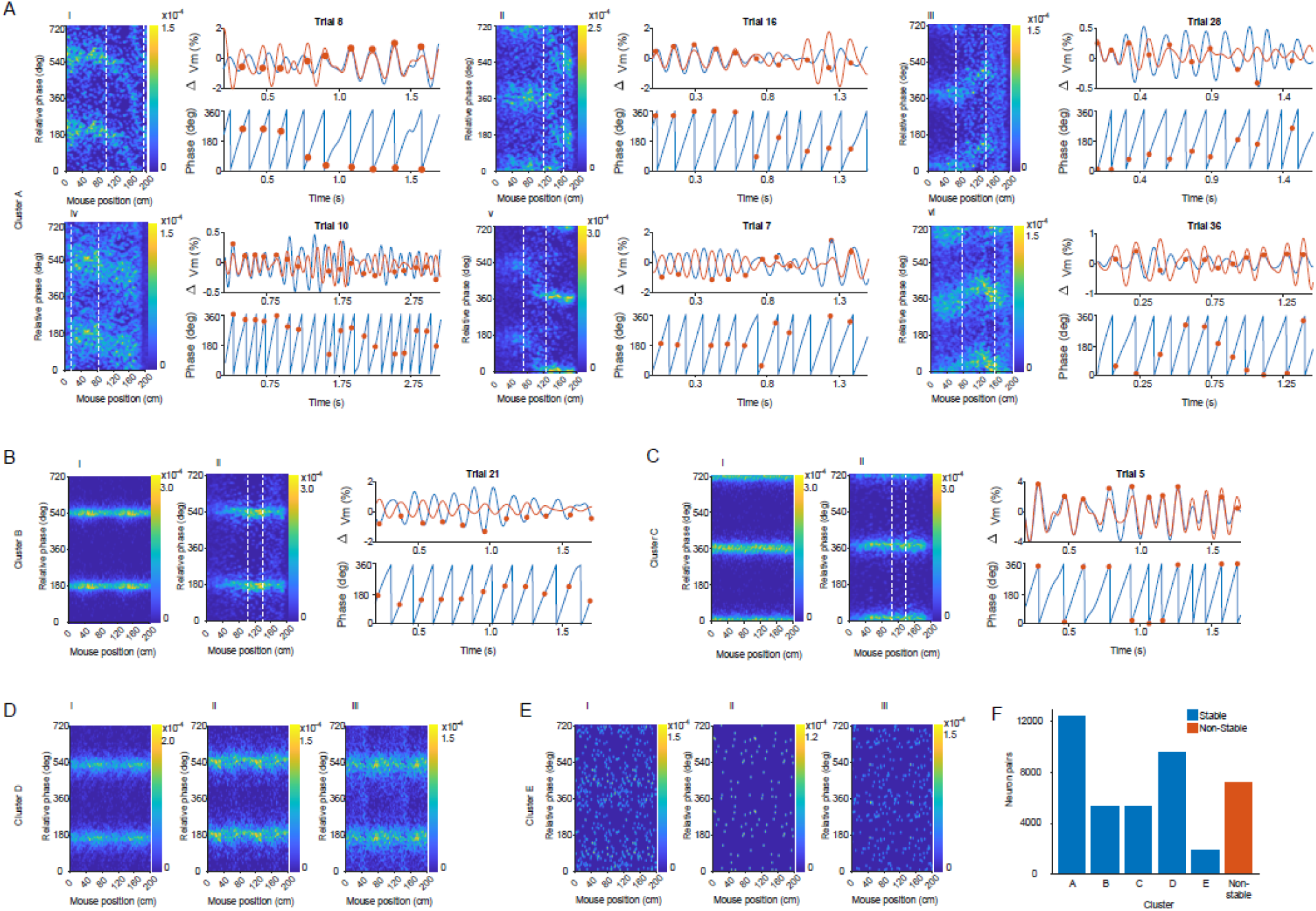
Relative shift of Vm theta phase between simultaneously recorded neuron pairs versus spatial position. (A-E). Examples of 5 distinct patterns of relative Vm theta phase shift of neuron pairs over position. Color indicates the probability of theta cycles at a given phase and spatial position (A) 6 example 2D histograms in cluster A showing gradual Vm theta phase shifts between neuron pairs over position. (i, iv) Phase precession, (ii, v) abrupt phase shifts, (iii) phase procession, and (vi) phase precession followed by procession. Left, 2D Gaussian smoothed histograms with yellow color indicating larger number of cycles at that relative phase. Shown are two theta cycles. Right top, theta filtered Vm traces of the neuron pair (orange and blue respectively) during a single representative traversal between the positions marked with dashed lines on the heatmap. Orange dots indicate the timing of theta peaks in the orange neuron. Right bottom, the theta peaks (orange dots) of the orange neuron overlaid on the instantaneous theta phase of the blue neuron. (B, C, D) Similar to A, but for example 2D histograms from clusters B, C and D respectively, showing consistent strong phase locking over positions. (E) Example 2D histograms from cluster E showing no obvious phase-position trends. (F) The number of neuron pairs in each cluster.

## DISCUSSION

The voltage imaging data from multiple simultaneously recorded neurons presented here introduce a novel framework for spatial coding in the hippocampus. This new framework shows coding of spatial location based on the relative phase dynamics of Vm oscillations between individual neurons. This coding by relative phase is independent of spiking, and can occur in neurons that are not spiking at all. The patterns of phase coding across different neuron pairs fall into multiple clusters. One cluster contains pairs of neurons that show systematic and gradual shifts in relative Vm theta phase during changes in virtual position (Figure 4Ai, iii, vi). This same cluster includes pairs of neurons that show abrupt shifts in relative phase of Vm from synchrony to antisynchrony at specific virtual positions, often around the reward delivery zone (start or end of the trial, Figure 4Aii, v). Finally, another two clusters contain pairs of neurons that maintain phase synchrony (Figure 4B, 4D) and one cluster has phase antisynchrony (Figure 4C) across all positions.

By performing simultaneous voltage imaging in multiple neurons, our results extend beyond earlier work using intracellular recording of single neurons in the hippocampus (Harvey et al., 2009) and entorhinal cortex (Domnisoru et al., 2013) that reported relative oscillation phase shifts of Vm versus LFP. In contrast, we show systematic shifts in Vm phase between neurons. By directly recording subthreshold Vm, our work also extends beyond previous extracellular recordings, demonstrating a shift of spiking from late to early phases of LFP theta, known as theta phase precession (O’Keefe & Recce, 1993; Skaggs et al., 1996; Kay et al., 2020). In contrast, our work reveals that the individual neuron’s Vm can have different relative phase relationships to other individual neuron’s Vm.

Most of the systematic phase shifts with position we observed are in cells without specific spiking firing fields, which is in general agreement with prior studies showing that spiking phase can code position even when there are no specific firing fields dependent on spatial location (Tingley & Buzsáki, 2018). In addition, the majority of the examples of relative phase are based on cycles without generation of spikes on that cycle. This coding scheme based on phase of subthreshold oscillations may be particularly relevant in conditions where spiking is sparse or suppressed, yet network dynamics persist. The flexibility of phase codes under different sensory and reward contingencies along the virtual linear track suggests a potential mechanism for continuous computations integrating allocentric and egocentric reference frames for guiding behavior.

Computational models have suggested that the relative phase of theta oscillations can play an important role in coding location. This includes the original models of theta phase precession based on interactions of subthreshold Vm oscillations in individual neurons relative to the phase of oscillations from network synaptic input (O’Keefe & Recce, 1993) or interneuron dynamics (Bose et al., 2000; Bose & Recce, 2001), or based on mechanisms involving differences in Vm dynamics on different portions of the dendritic tree (Magee, 2001). Future studies with dendritic voltage imaging would reveal additional insights on the significance of how network inputs shape the observed somatic Vm phase shifts between neurons.

The relative theta phase of the Vm dynamics in hippocampal neurons could be regulated by input from the medial septum (Robinson et al., 2023, 2024). Neurons in the medial septum show a range of theta phase preferences (King et al., 1998), which could regulate the theta phase of hippocampal interneurons (Tóth et al., 1997; Hasselmo & Shay, 2014). The relative phase of Vm dynamics in hippocampus could also be influenced by input from the entorhinal cortex, as entorhinal neurons show strong phase coding relative to theta rhythm (Hafting et al., 2008; Brandon et al., 2013; Domnisoru et al., 2013) and lesions of entorhinal cortex impair theta phase precession in hippocampus (Mankin et al., 2012). Computational models show that theta can enhance the context dependent retrieval of sequences on specific phases of theta (Hasselmo, 2005; Hasselmo & Eichenbaum, 2005). On a functional level, the regulation of theta dependent phase coding could contribute to the phase dependent activation of future trajectories at specific time points (Johnson & Redish, 2007; Pfeiffer & Foster, 2013) or on alternating theta cycles (Brandon et al., 2013; Kay et al., 2020; Vollan et al., 2025). These theta dependent retrieval dynamics could be useful for guiding behavior based on overlap of future trajectories with reward locations (Erdem & Hasselmo, 2012).

The simultaneous voltage imaging of Vm in multiple different neurons provides an opportunity to test for new dimensions of neural coding at the subthreshold level. A fundamental question in neuroscience is how neurons code information in subthreshold Vm that is influenced not only by synaptic inputs but by intrinsic voltage dependent conductances. Is information coded by the mean depolarization over bins of time, or by complex temporal dynamics such as the relative phase of Vm oscillations between different neurons? Most abstract network models (e.g. Hopfield nets, recurrent neural networks) use mean magnitude of depolarization over time bins that generate mean firing rates. Mean firing rate has been used in this way in most attractor networks or Hopfield networks (Hopfield, 1984), as well as deep learning networks (LeCun et al., 2015) and recurrent neural networks (DePasquale et al., 2018). However, the importance of temporal dynamics is supported by the wide range of voltage-sensitive membrane conductances with broad time courses (Koch & Segev, 2000; Izhikevich, 2006). Because most in vivo data in behaving animals uses extracellular recording, there has been a focus on coding by the spiking of neurons. However, the activation of spikes at discrete time points might reflect only intermittent sampling of more continuous temporal coding by the relative phase of subthreshold dynamics of activated cells that is intermittently accessed by generation of spikes for the transfer of information. This highlights the importance of being able to analyze the relative phase of membrane potential Vm oscillations or other dynamics of membrane potential across multiple simultaneously recorded neurons relative to position.

## METHODS

### Animal preparation and habituation

Experiments were performed in 7 adult C57BL/6 mice, both male and female. Animals were first habituated to handling for > 3 days prior to surgery. During aseptic surgery, a custom imaging window, coupled with an infusion cannula and an LFP electrode was implanted over the CA1 region (center of window coordinates: AP: -2.00 mm, ML: +2.00 mm, DV: -1.6 mm depth) after the gentle removal of the overlaying cortex. A head bar was also attached to the skull during the same surgery. After recovery from surgery (> 7 days), a mixture of 500nL of AAV1-Syn-FLEX-Voltron2-WPRE (>5x10^12 GC/ml, Addgene, catlog #: 172907) and 500nL of AAV9-CamKII-Cre (2.1x10^13 GC/ml, Addgene, catlog #: 182736) was infused through the infusion cannula to express Voltron2 in CA1. At the concentration used, Voltron2 was expressed in CA1 neurons nonselectively, as also confirmed by histology (Supplementary Figure 7).

### Virtual spatial task

We used a virtual spatial navigation task similar to that in Pettit et al., 2022. Specifically, water-restricted mice were head fixed under the microscope and they voluntarily moved on a spherical treadmill in a virtual environment executed in MATLAB with ViRMEn (Aronov & Tank, 2014). The spherical treadmill was supported by an axle that ran through the ball’s diameter. Mice were trained to run unidirectionally on a continuously looped linear track. Each lap was 200 cm long, containing distinct proximal and distal virtual visual cues (shapes, patterns, and colors), and a water reward zone at the end.

Animals were trained for the task in 4 stages. In stage 1, the animals were placed on a Styrofoam ball, supported by air, allowing movement in any direction to habituate to head-fixation. In stage 2, mice ran on the unidirectional spherical treadmill during each training session of 30 minutes. During the first two minutes of the session, 1uL of water was provided at every 10s interval to introduce the water lick port. During the subsequent 28 minutes, water reward was provided if the mouse performed either a forward movement or licking. In stage 3, we introduced the animals to the virtual world on the spherical treadmill, in addition to the stage 2 water rewards. In Stage 4, water reward was provided only at the reward zone when the mouse completed a traversal (trial) in the virtual world. On “standard trials,” the mice received water reward if it licked the water port while inside the reward zone. On the rest of trials (“crutch trials”), reward was administered regardless of licking behavior to help mice learn the reward zone. Number of crutch trials started at 95% and was gradually decreased to 15% as the mouse learned the task. Once mice reached the performance criteria (described in section on behavior analysis) we started voltage imaging.

### Movement data acquisition

Movement of the ball was tracked using a computer mouse sensor (Glorious Model D Gaming Mouse) placed at the equator of the Styrofoam ball. The mouse sensor’s y-surface displacement data was read using an Arduino Uno and a custom Arduino script, scaled to match the actual speed of the ball rotation in cm/sec (using a calibration scale factor determined by moving the ball at a known speed), and fed into VirMEn to update the virtual world.

### Lick data acquisition and reward delivery

Licks were detected in real-time through tracking the mouse tongue with DeepLabCut-Live (Kane et al., 2020). The DeepLabCut-Live implementation used a pretrained and validated machine learning algorithm to detect licking using an incoming video stream from a webcam (Logitech-4K-Pro-Webcam). When the tongue was detected in a video frame, a custom Python function sent control signals via an Arduino device to the VirMEn computer to indicate a lick. Rewards were administered using TTL pulses generated by a DAQ board (USB-6259, National Instruments) to turn on a solenoid pump to deliver water.

### LFP recording

LFPs were recorded via custom electrodes, a head stage and an LFP amplifier (Tucker Davis Technologies, PZ2) using the recording software Synapse at 2kHz sampling rate.

### Behavior analysis

To evaluate the performance of the VR task, we used previously published lick selectivity metric (Pettit et al., 2022). In the linear VR track, an area of 25cm marked with unique floor patterns was defined as the reward zone. A zone of equal size at the midpoint between two rewards was defined as the opposite zone. Anticipatory licks (licks prior to when the reward is administered) were counted in both the reward zone and the opposite zone. Lick selectivity was defined as *lick selectivity = (reward zone licks - opposite zone licks) / (reward zone licks + opposite zone licks).* The criterion for inclusion was lick selectivity > 0.6 at the end of training stage 3.

### Voltron2 voltage imaging data collection

A genetically encoded voltage indicator, Voltron2, was used. At least one day before voltage imaging, an aliquot of JF_552_-HaloTag ligand (100nmol) was dissolved into a solution consisting of anhydrous DMSO (20uL), 20% (w/v) Pluronic F-127, and PBS (80uL). This solution was retro-orbitally administered following the procedure described in Abdelfattah et al., 2023. Each JF_552_-HaloTag ligand injection allows Voltron2 labelled cells to be imaged for up to 5 days before the need of re-administering the dye. During each recording session, animals were positioned under a custom digital micromirro device (AJD-4500, Ajile Light Indusries) based targeted illumination microscope (Xiao et al., 2021), and performed the virtual spatial task described above. The microscope was equipped with a scientific complementary metal oxide semiconductor (sCMOS) camera (Kinetix, Teledyne Photometric) and a 40x objective (Nikon, 40×/0.8NA CFI APO NIR). Voltron2 was excited by a 561 nm 100mW laser (Cobolt 06-DPL, HÜBNER Photonics). Images were acquired using Beacon/PVCam software (Teledyne Photometric) at 800Hz, with each frame capture triggered by TTL pulses generated by a DAQ board (USB-6259, National Instruments). TTL pulses were recorded using a RZ5D recording system (Tucker Davis Technologies).

### Voltage imaging data preprocessing and Vm trace extraction

With targeted illumination, each full field of view contained many illuminated areas. We first manually segmented each illuminated area using a max-min projection image. Each region was then separately motion corrected using a custom adaptation of the NoRMCorre algorithm (Pnevmatikakis & Giovannucci, 2017), in two steps. First, the video frames were divided into non-overlapping subsets of 10000 consecutive frames, and each set was motion-corrected independently. Next, the 2D projections of each set of frames were motion corrected. The corrective shifts calculated from the two steps in both X, and Y direction were then combined and applied to the raw videos to obtain the motion corrected videos.

A max-min projection image created from the motion-corrected videos was then used for manual cell segmentation. Fluorescence for each cell was then obtained as the average fluorescence intensity *F*(*t*) across all pixels in that cell minus camera dark noise measured with no illumination light. A baseline fluorescence trace, *F*_0_(*t*) was computed by filtering with a 1s moving average. Next, to correct for photobleaching, and to invert the decrease in Voltron fluorescence during depolarization, the fractional change 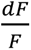 in fluorescence was calculated as 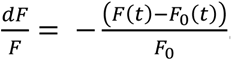. The resulting trace was considered the Vm of a given neuron.

### Image motion rejection

To identify out-of-focus frames (with Z-direction image motion), we developed a focus measurement algorithm. First, we manually selected a rectangular ROI around each neuron of the motion corrected image stack, and computed the 2D FFT for each image frame. We then binned the 2D FFT spatial frequencies into 30 bins and computed the corresponding power to generate a feature vector for each image frame.

We then examined each 2-second-long recording segment (1600 frames) to determine if the recording segment contained in-focus versus out-of-focus frames. To do so, we performed a cluster separation analysis. Specifically, we clustered the spatial frequency feature vectors generated from the 1600 frames into 2 clusters using a Gaussian mixture model (Matlab function ‘fitgmdist’). To evaluate the separation of the 2 clusters, we calculated the silhouette scores *s*(*i*) for each frame *i* using equation 1.

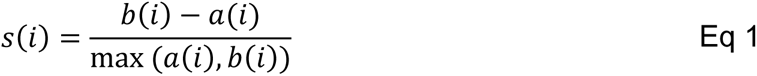

Where, *a*(*i*) is the average distance from frame *i* to all other frames in the same cluster, and *b*(*i*) is the average distance from frame *i* to all other frames in the other cluster.

If the mode (peak) of the distribution of the silhouette scores was greater than 0.6, the 2 clusters were deemed well separated, containing frames with two different focus levels. We created a binary cluster identity vector of the same length as the number of frames in the segment (1600) and assigned 0s or 1s to each frame indicating their cluster identity. Since Z motion corresponds to the shift in focus, we identified the frames in which the focus transitions occurred (image motion onset) by computing the standard deviation of the cluster identity vector using a moving window of 100ms, and thresholding at 0.5 standard deviations. To examine any image motion at the junction of 2-second-long recording segments, in the 2^nd^ iteration, we repeated the cluster separation analysis, but considering a new set of 2-second-long recording segments centered on the junction of the 1^st^ iteration. The union of the two iterations were used to identify all potential frames with Z-motion.

Using the same procedure, we identified potential frames with Z-motion for all neurons within a recording. Because Z-motion is expected to impact all the neurons within a recording, to determine whether there is Z-motion at each time point, we used a threshold where the number of neurons with potential Z-motion must be greater than 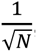, where *N* is the number of neurons in the recording.

To identify frames with X or Y-direction image motion, we identified Vm fluctuations that are correlated to the corrective shifts applied by the motion correction algorithm. First, we obtained a regression fit for Vm as a function of the corrective shifts using Equation 2.

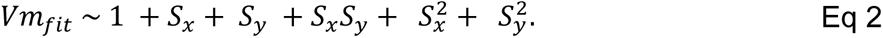

Where, *Vm_fit_* is the regression fit for *V_m_*, and *S_x_* and *S_y_* are the corrective shifts applied to each imaging frame in the x and y direction respectively. We then smoothed *Vm* and *Vm_fit_* using a moving average filter of width 10ms and computed a correlation trace *f_corr_* the resulting smoothed *Vm* and *Vm_fit_* traces with a moving window of 200ms.

Consecutive time points with *f*_*corr* > 0.75 and 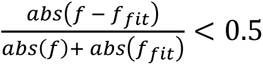 for >500ms were identified as candidate windows with X or Y-direction motion. We then computed the moving Vm mean of 10ms, and identified frames with X-Y motion as those with Vm above the 99^th^ percentile in a 3s neighborhood. Frames identified as having X-Y or Z motion were excluded from all subsequent analyses.

### Spike detection

Spikes were detected using a previously published method with some modifications (Abdelfattah et al., 2023). Briefly, the Vm trace was high-pass filtered by subtracting a median filtered trace. Next, all positive peaks of this trace were detected as candidate spikes. To determine a threshold, the high-pass filtered trace was inverted (to obtain a ‘noise’ trace), and positive peaks of this trace were detected similarly. Next, two probability distributions were built for the amplitudes of these two sets of peaks *ps* (*x*) and *pn* (*x*) respectively. The ‘spike’ amplitude distribution is computed as *pk* (*x*) = *ps* (*x*) − *pn* (*x*). An amplitude threshold *t* was determined such that *pk* (*t*) = *pn* (*x*). Next, a false positive rate (FPR) is determined by applying this threshold to the inverted trace, and counting the number of spikes detected. If FPR was greater than 0.1 Hz, then the threshold was increased until the FPR drops to 0.1 Hz. This final threshold was then applied to the high-pass filtered signal to determine spike timing.

### Phase-position analysis

For phase-position analysis, we first filtered Vm with an FIR bandpass filter at the theta frequencies (6-10Hz), and obtained the instantaneous phase as the angle of the Hilbert transform of the filtered Vm. Peaks of each theta cycle corresponded to 0 instantaneous phase. Time periods when mouse movement speed was less than 10 cm/s were excluded from subsequent analyses. To analyze the relationship between the relative Vm theta phase of neuron pairs, and the mouse position, we first identified the peaks of each theta cycle in the first neuron, and the corresponding spatial location of the mouse at that time. For each identified theta peak, the instantaneous theta phase of the second neuron was defined as the relative phase between the neuron pair.

We then created a 2D histogram by binning the relative phases and the corresponding mouse positions into 1-degree x 1 cm bins. The resulting histogram was normalized by the occupancy in each position bin, and smoothed using a 2D - Gaussian filter with a width 9 bins and a standard deviation of 2 bins.

To evaluate the stability of the phase-position histograms, we generated two separate histograms using either the first half of the trials or the second half of the trials, and computed Spearman’s correlation of the two histograms. We then obtained a shuffled distribution of the Spearman’s correlation coefficient by repeating this procedure 1000 times using shuffled Vm traces by circularly shifting the two traces by a random duration. If the actual Spearman’s correlation coefficient was higher than the 95^th^ percentile of the shuffled distribution, it was considered a stable histogram.

To cluster 2D histograms, each stable position-phase histogram was converted to an 180x180 pixel greyscale image, and normalized between 0 and 1. We then trained an autoencoder of hidden neuron size 1000 to encode and decode these images. The trained autoencoder was used to encode each input image by a sparse feature vector of length 1000. The histograms were then clustered using this feature vector and k-means clustering to obtain 5 clusters.

## Acknowledgements

1. X. H. acknowledges funding from NIH 1R01MH122971 and NSF 2002971-DIOS. X. H. and M. E.H. acknowledge funding from Boston University Kilachand award. S. X. and J. M. acknowledge support from NIH R34NS127098. C.R. acknowledges support from NSF GRFP2021324226. D.S. acknowledges funding from the American Epilepsy Foundation Predoctoral Fellowship. This work was additionally supported by Boston University Micro and Nano Imaging Facility through NIH S10OD024993.

## Authorship contribution statement

Conceptualization: MA, XH, MEH, SM, RM

Data collection & analysis: MA, SM, RM

Experimental support: RM, KP, YW, DS, CR, ER

Reagent, material and data analysis support: SX, HT, SEP, JBG, LDL, JM, DS

Writing – Original Draft: MA, SM, XH, MEH

Writing – Review & Editing: All authors Supervision: XH, MEH

## Supplementary Material

**Supplementary Table 1:**
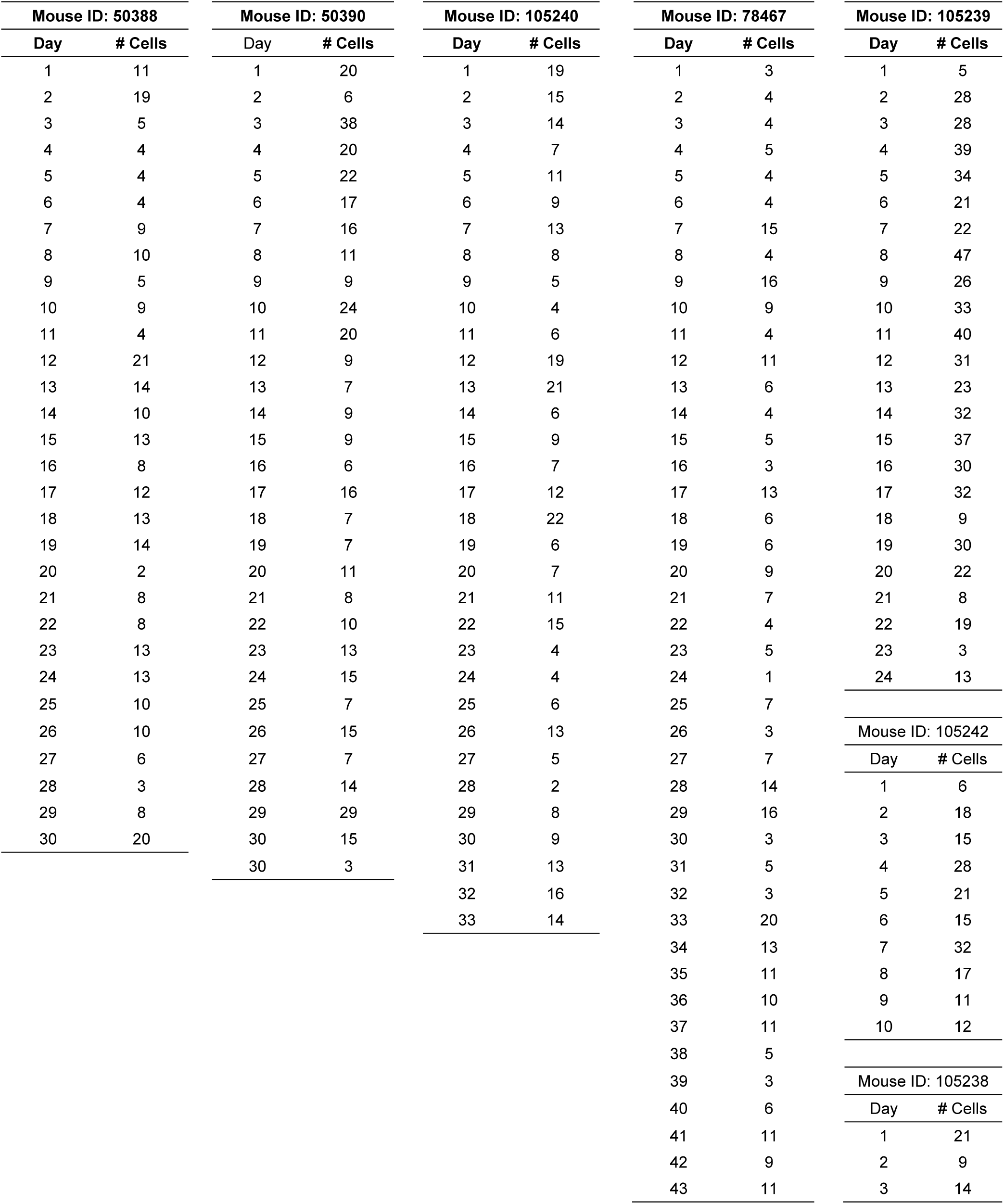
Number of neurons recorded in each session.

**Supplementary Figure 1:**
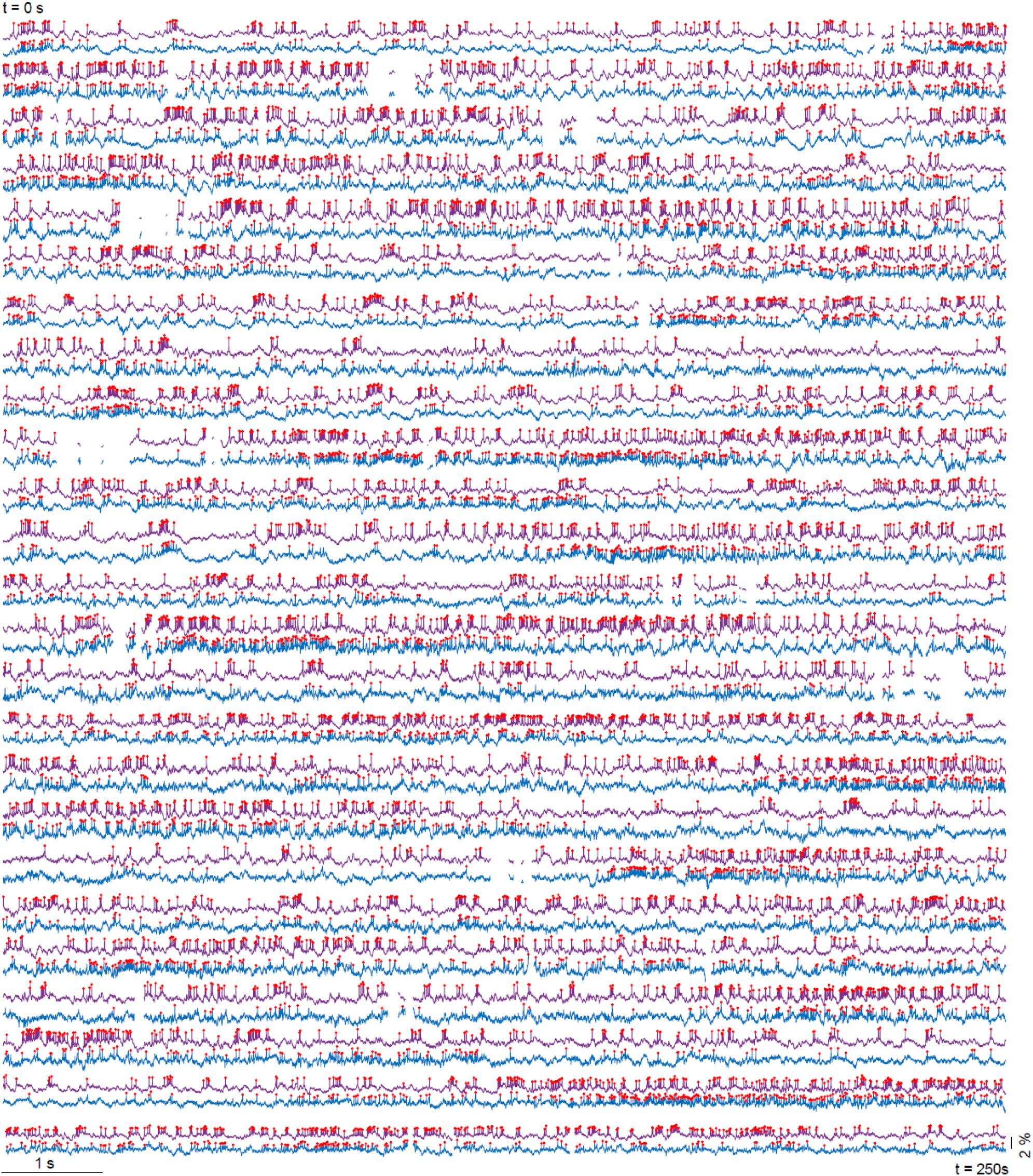
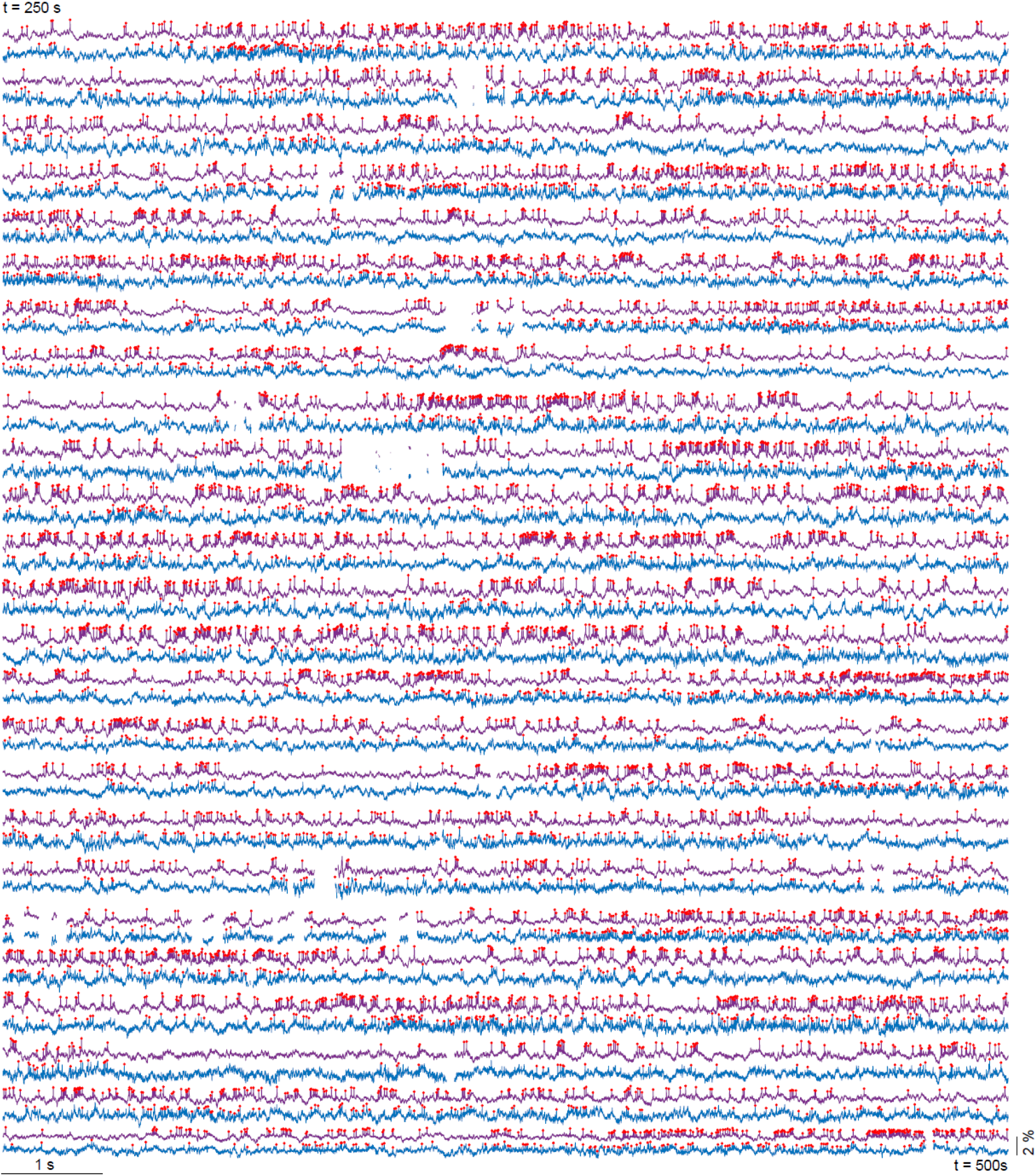
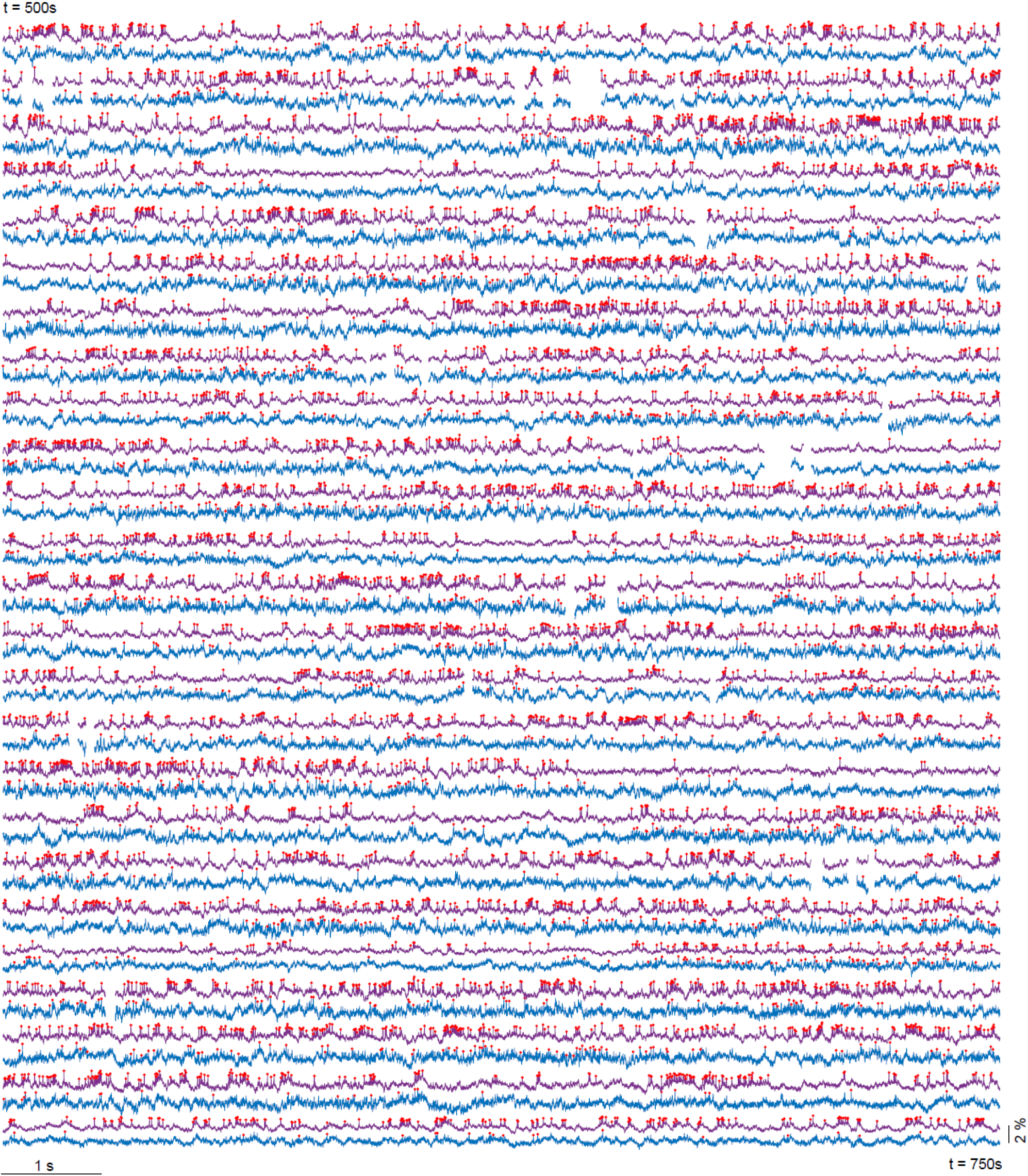
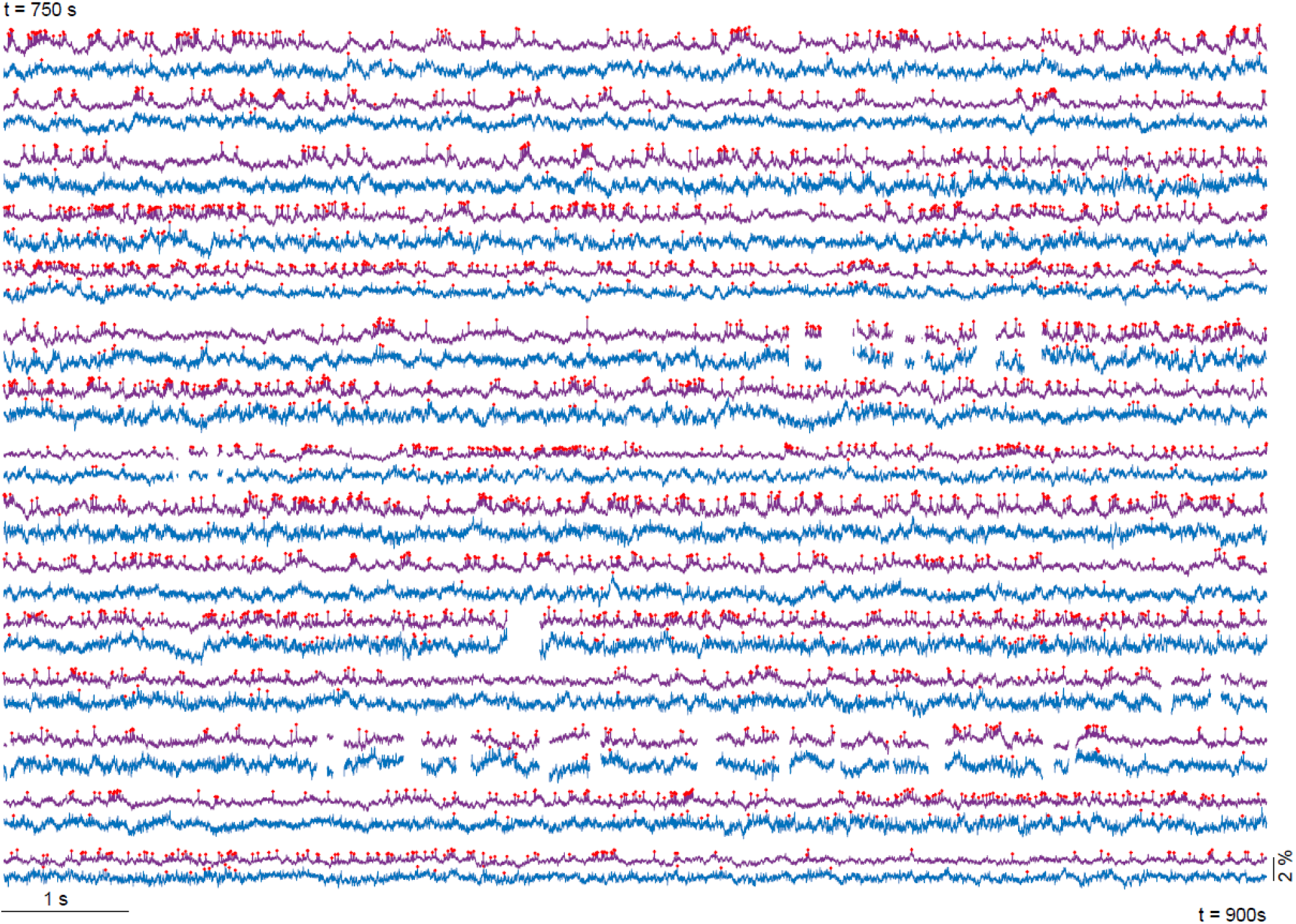
An example 15-minute recording session with 2 simultaneously recorded neurons. Vm traces of the neurons are shown in blue and purple respectively. Red asterisks denote identified spikes. Time points with significant image motion were excluded.

**Supplementary Figure 2:**
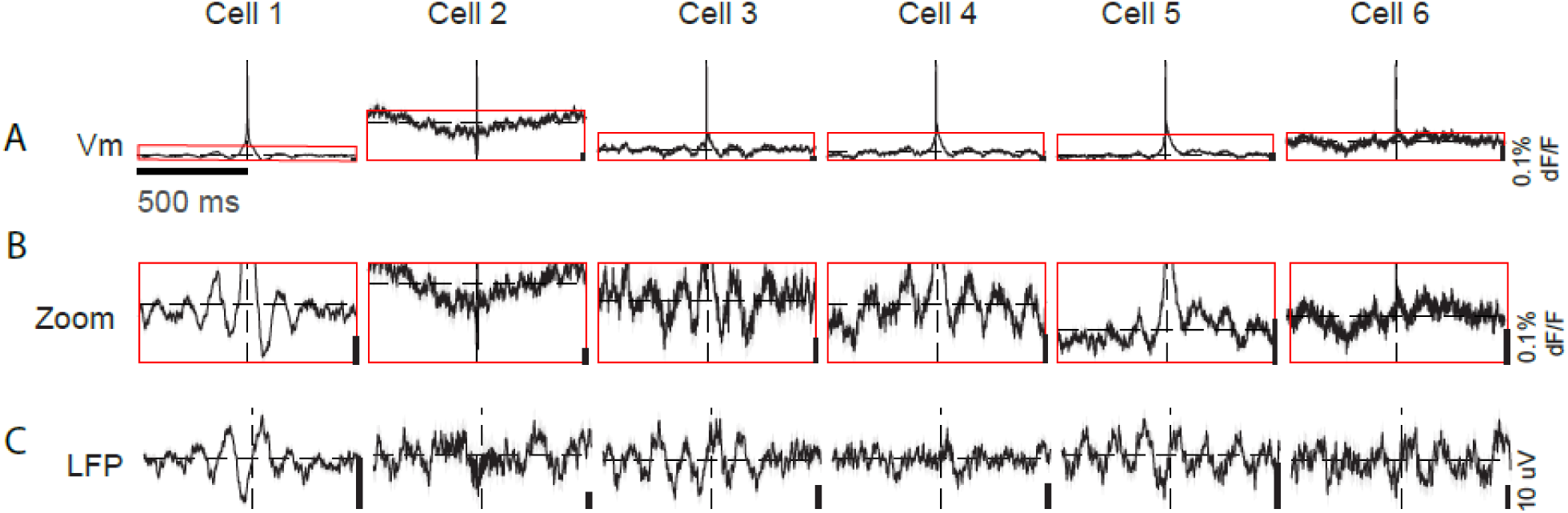
**Spiking predominantly occurred on the rising phase of Vm and LFP theta oscillations**. An example session showing (A) Vm of a neuron aligned to the identified spikes. (B) Zoom-ins of A. (C) LFP aligned to the identified spikes for each neuron. Black lines are mean, and shaded areas are SEM. Dashed lines indicate 0.

**Supplementary Figure 3:**
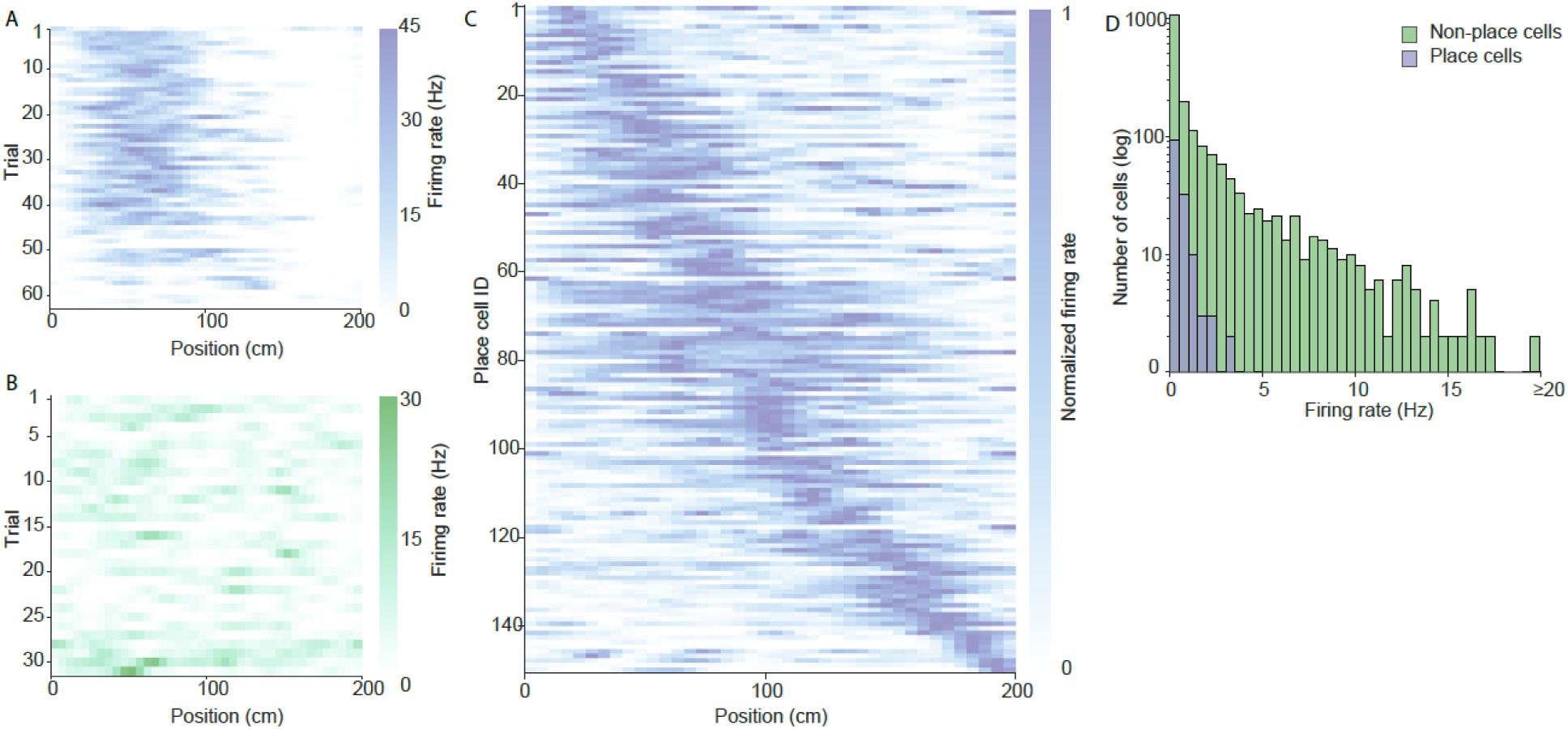
Place cells showing spatially specific spiking. Example rate map of (A) a place cell and (B) a non-place cell. (C) The identified place cells tiled the entire virtual track. Neurons were sorted by the location of their place fields. Only the neurons containing a single place field are included (n = 7 mice, 143 neurons). (D) Firing rate distribution of place cells and non-place cells.

**Supplementary Figure 4:**
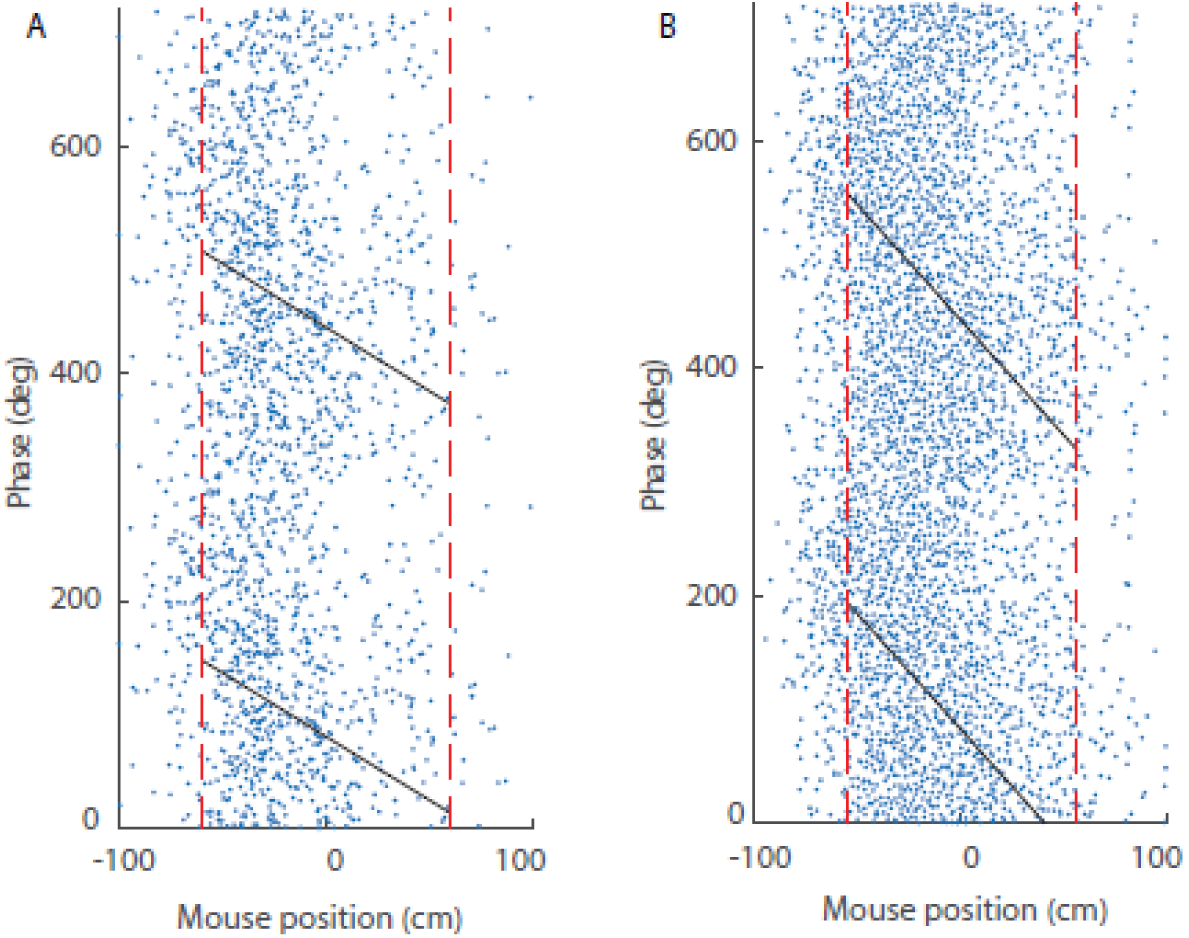
Example phase precession in two simultaneously recorded place cells. (A) Spike timing relative to LFP theta phase. Black lines are the best fit linear-circular regression. The vertical red dashed lines mark the beginning and end of the place field. X-axis is centered to the place field. Y-axis shows two theta cycles. B) Same as in A, for a different neuron.

**Supplementary Figure 5:**
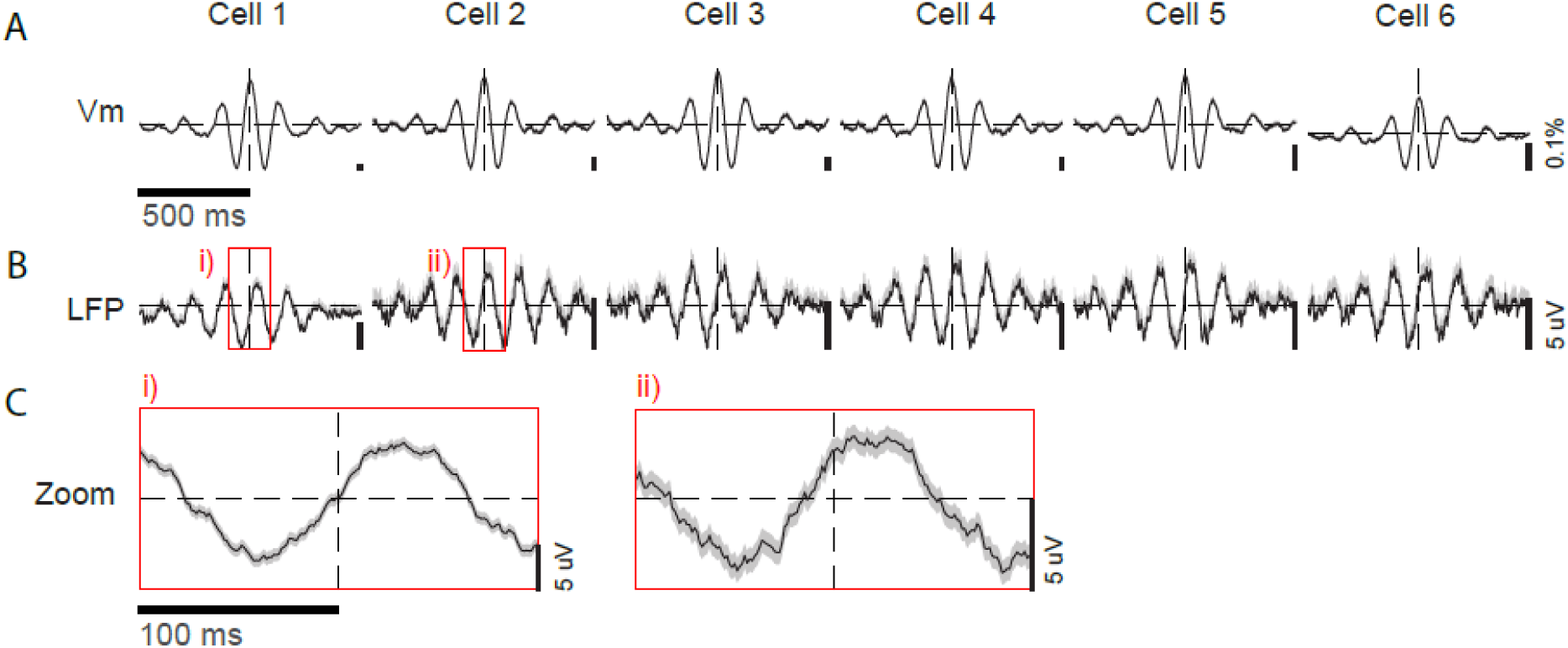
**Vm theta peak in many cells occurred at specific phases of LFP theta**. (A) An example session showing Vm of simultaneously recorded neurons aligned to the peak of their own Vm theta. (B) LFP aligned to the identified spikes of each neuron. (C) Zoom-ins of B. Black lines are mean, and shaded areas are SEM. Dashed lines indicate 0.

**Supplementary Figure 6:**
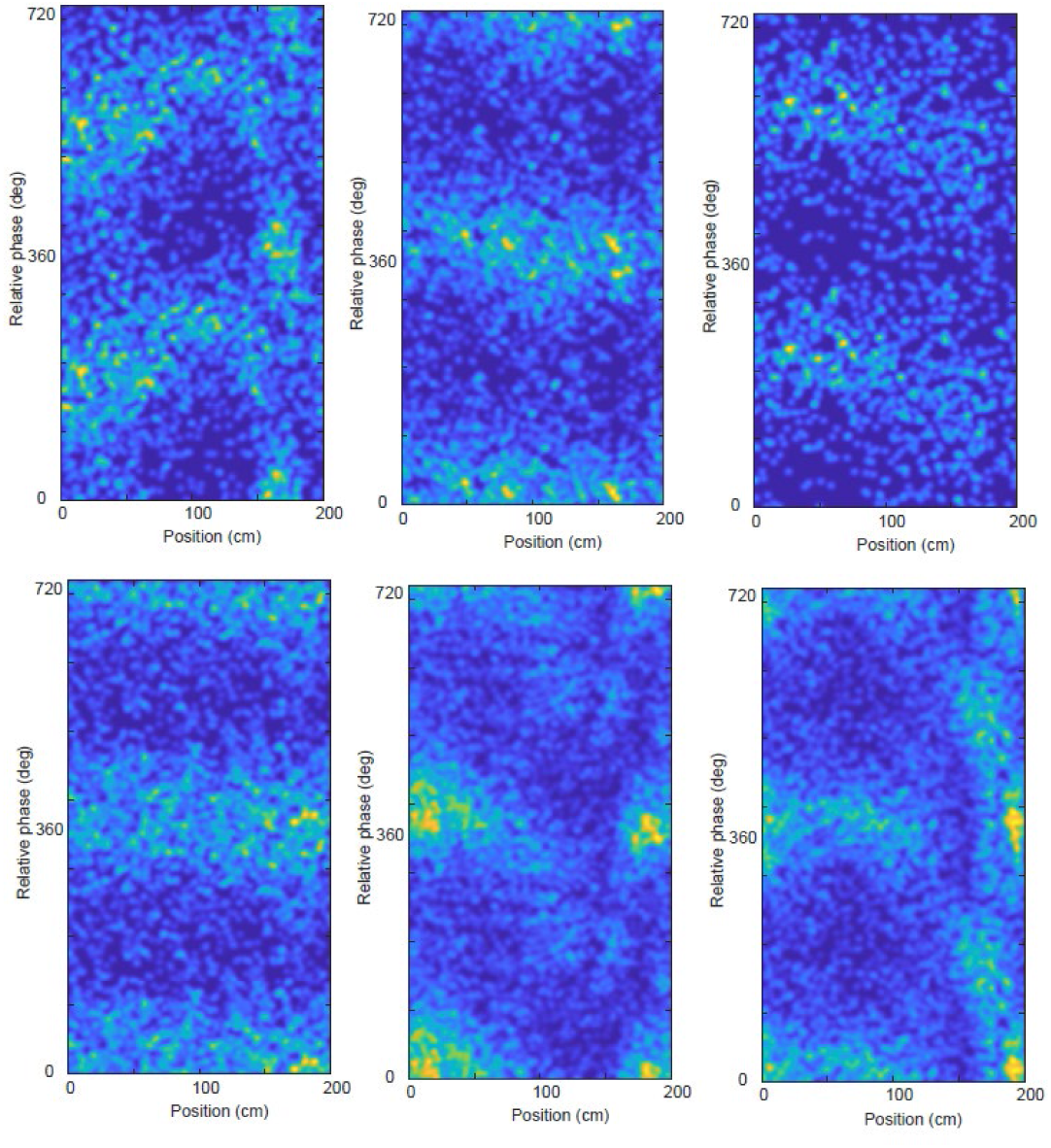
Additional examples of the **relative phase of neuron pairs in Cluster A that dynamically changed along with spatial position**. Six additional example 2D histogram heatmaps for neuron pairs exhibiting gradual shifting of their relative phase over spatial position. These are additional examples from the same cluster as the six cell pairs shown in Figure 4A.

**Supplementary Figure 7:**
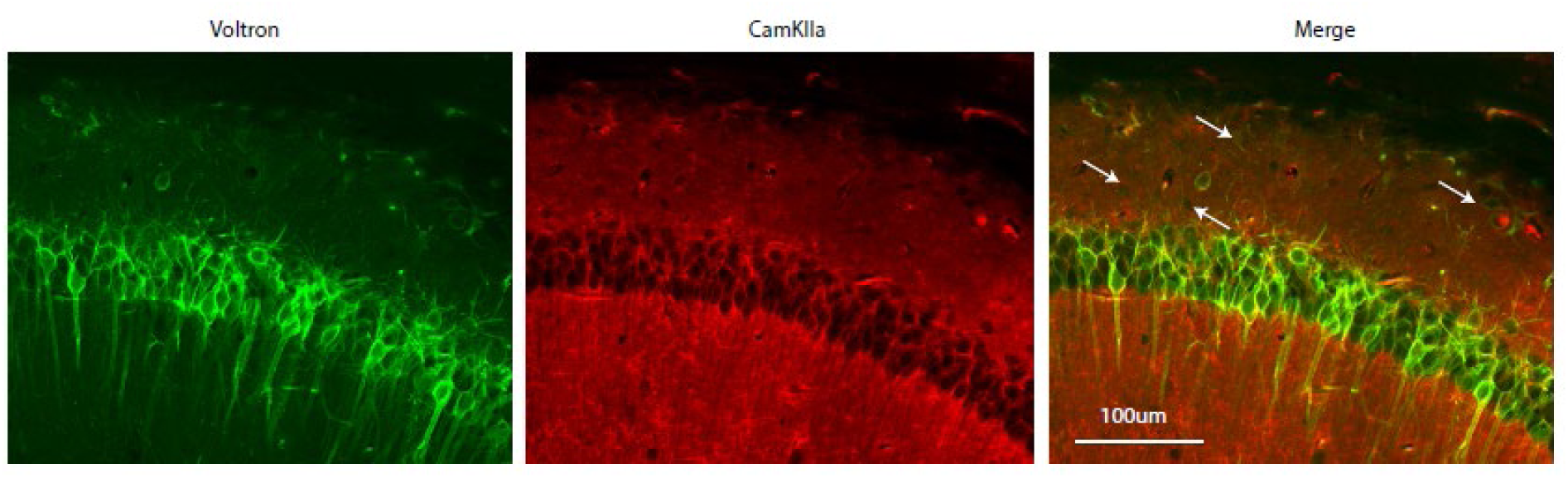
**Voltron2 non-specifically labeled CA1 neurons**. Left, a representative confocal image showing Voltron 2 expression (Green). Middle, immunofluorescence of CamKIIa (red). Right, merge. Arrows indicate cells in the Stratum Oriens, that are positive or voltron2 but not for CamKIIa immunofluorescence.

## Notes

### Competing Interest Statement

The authors have declared no competing interest.

