## Supplemental Table and Figures for "Relative phase of membrane potential theta oscillations between individual hippocampal neurons code space"

### Supplementary Material

### Supplementary Table 1: Number of neurons recorded in each session

| Mouse ID: 50388 | |
|---|---|
| **Day** | **# Cells** |
| 1 | 11 |
| 2 | 19 |
| 3 | 5 |
| 4 | 4 |
| 5 | 4 |
| 6 | 4 |
| 7 | 9 |
| 8 | 10 |
| 9 | 5 |
| 10 | 9 |
| 11 | 4 |
| 12 | 21 |
| 13 | 14 |
| 14 | 10 |
| 15 | 13 |
| 16 | 8 |
| 17 | 12 |
| 18 | 13 |
| 19 | 14 |
| 20 | 2 |
| 21 | 8 |
| 22 | 8 |
| 23 | 13 |
| 24 | 13 |
| 25 | 10 |
| 26 | 10 |
| 27 | 6 |
| 28 | 3 |
| 29 | 8 |
| 30 | 20 |

| Mouse ID: 50390 | |
|---|---|
| Day | **# Cells** |
| 1 | 20 |
| 2 | 6 |
| 3 | 38 |
| 4 | 20 |
| 5 | 22 |
| 6 | 17 |
| 7 | 16 |
| 8 | 11 |
| 9 | 9 |
| 10 | 24 |
| 11 | 20 |
| 12 | 9 |
| 13 | 7 |
| 14 | 9 |
| 15 | 9 |
| 16 | 6 |
| 17 | 16 |
| 18 | 7 |
| 19 | 7 |
| 20 | 11 |
| 21 | 8 |
| 22 | 10 |
| 23 | 13 |
| 24 | 15 |
| 25 | 7 |
| 26 | 15 |
| 27 | 7 |
| 28 | 14 |
| 29 | 29 |
| 30 | 15 |
| 30 | 3 |

| Mouse ID: 105240 | |
|---|---|
| **Day** | **# Cells** |
| 1 | 19 |
| 2 | 15 |
| 3 | 14 |
| 4 | 7 |
| 5 | 11 |
| 6 | 9 |
| 7 | 13 |
| 8 | 8 |
| 9 | 5 |
| 10 | 4 |
| 11 | 6 |
| 12 | 19 |
| 13 | 21 |
| 14 | 6 |
| 15 | 9 |
| 16 | 7 |
| 17 | 12 |
| 18 | 22 |
| 19 | 6 |
| 20 | 7 |
| 21 | 11 |
| 22 | 15 |
| 23 | 4 |
| 24 | 4 |
| 25 | 6 |
| 26 | 13 |
| 27 | 5 |
| 28 | 2 |
| 29 | 8 |
| 30 | 9 |
| 31 | 13 |
| 32 | 16 |
| 33 | 14 |

| Mouse ID: 78467 | |
|---|---|
| **Day** | **# Cells** |
| 1 | 3 |
| 2 | 4 |
| 3 | 4 |
| 4 | 5 |
| 5 | 4 |
| 6 | 4 |
| 7 | 15 |
| 8 | 4 |
| 9 | 16 |
| 10 | 9 |
| 11 | 4 |
| 12 | 11 |
| 13 | 6 |
| 14 | 4 |
| 15 | 5 |
| 16 | 3 |
| 17 | 13 |
| 18 | 6 |
| 19 | 6 |
| 20 | 9 |
| 21 | 7 |
| 22 | 4 |
| 23 | 5 |
| 24 | 1 |
| 25 | 7 |
| 26 | 3 |
| 27 | 7 |
| 28 | 14 |
| 29 | 16 |
| 30 | 3 |
| 31 | 5 |
| 32 | 3 |
| 33 | 20 |
| 34 | 13 |
| 35 | 11 |
| 36 | 10 |
| 37 | 11 |
| 38 | 5 |
| 39 | 3 |
| 40 | 6 |
| 41 | 11 |
| 42 | 9 |
| 43 | 11 |

| Mouse ID: 105239 | |
|---|---|
| **Day** | **# Cells** |
| 1 | 5 |
| 2 | 28 |
| 3 | 28 |
| 4 | 39 |
| 5 | 34 |
| 6 | 21 |
| 7 | 22 |
| 8 | 47 |
| 9 | 26 |
| 10 | 33 |
| 11 | 40 |
| 12 | 31 |
| 13 | 23 |
| 14 | 32 |
| 15 | 37 |
| 16 | 30 |
| 17 | 32 |
| 18 | 9 |
| 19 | 30 |
| 20 | 22 |
| 21 | 8 |
| 22 | 19 |
| 23 | 3 |
| 24 | 13 |

| Mouse ID: 105242 | |
|---|---|
| Day | # Cells |
| 1 | 6 |
| 2 | 18 |
| 3 | 15 |
| 4 | 28 |
| 5 | 21 |
| 6 | 15 |
| 7 | 32 |
| 8 | 17 |
| 9 | 11 |
| 10 | 12 |

| Mouse ID: 105238 | |
|---|---|
| Day | # Cells |
| 1 | 21 |
| 2 | 9 |
| 3 | 14 |

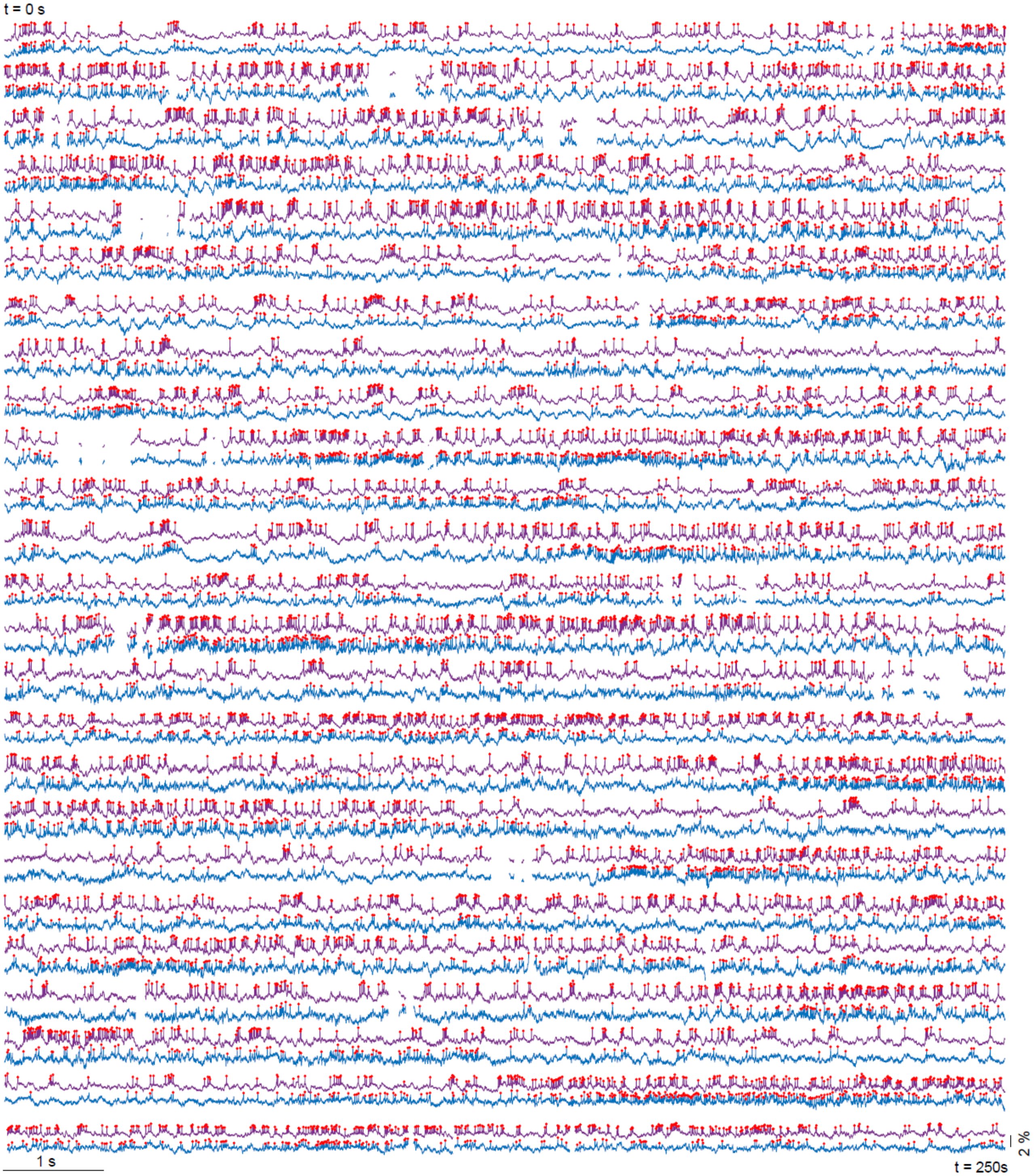
t = 0 s
1 s
2 %
t = 250s

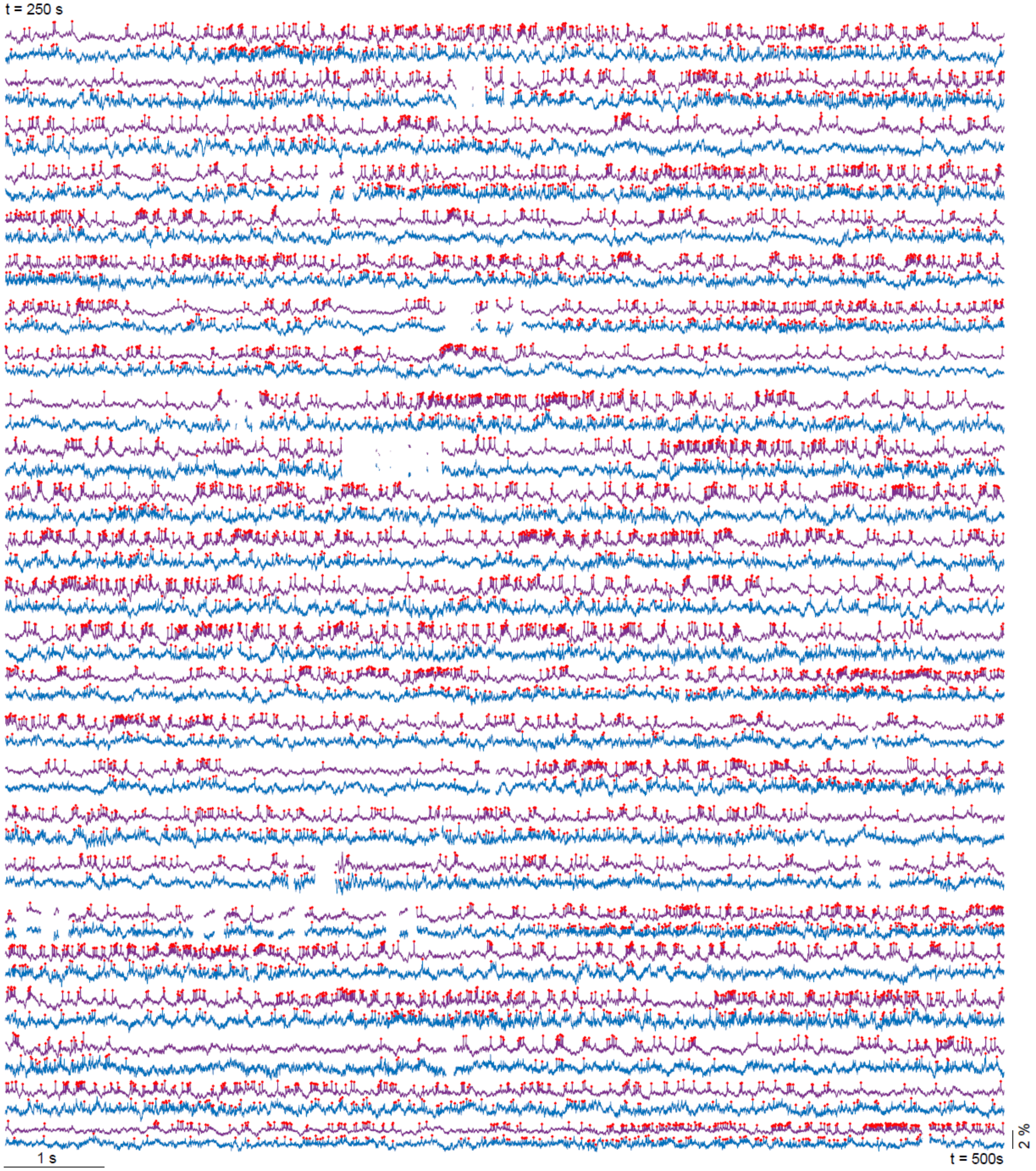
t = 250 s
1 s
t = 500s
2 %

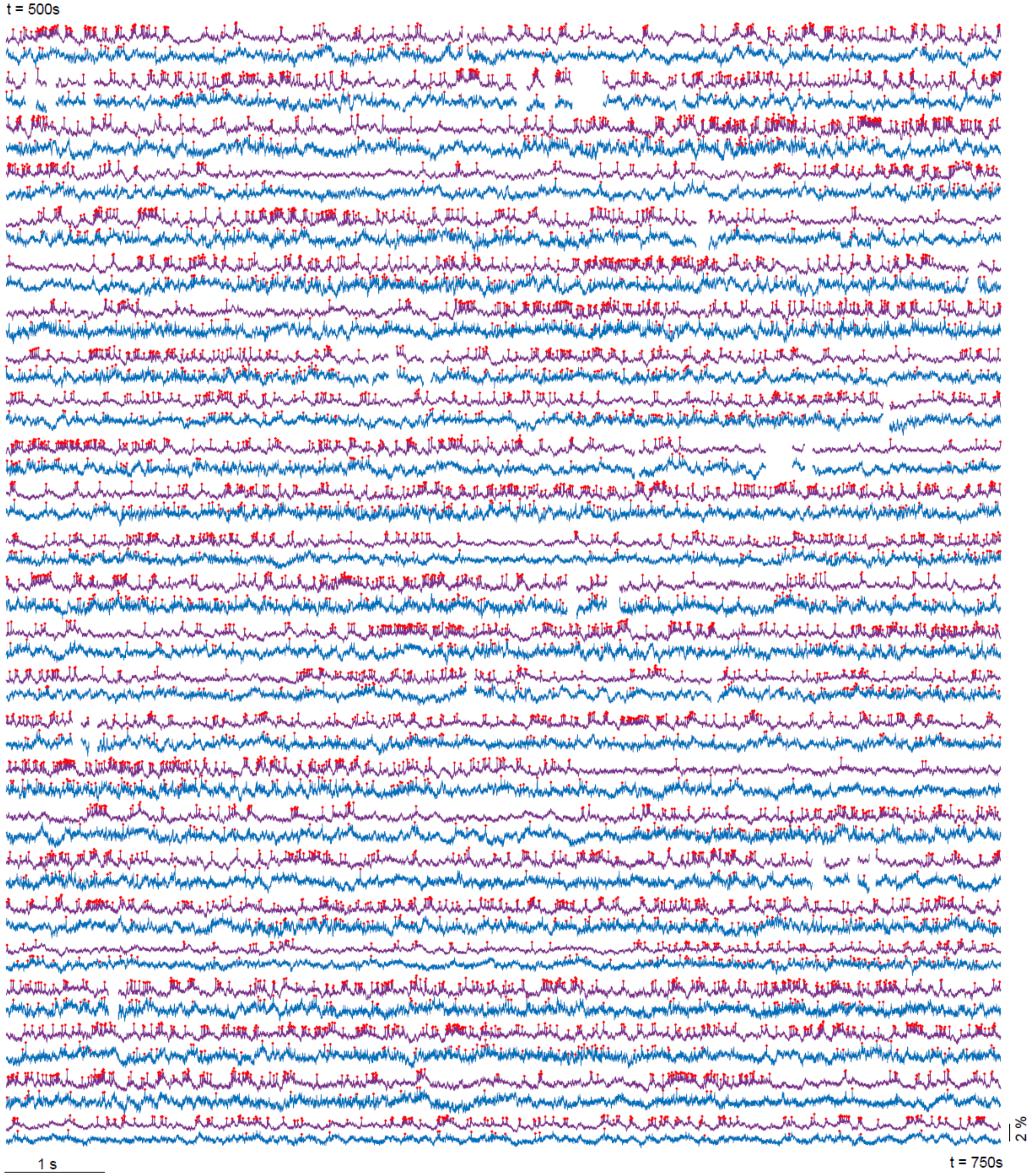
t = 500s
1 s
t = 750s
2 %

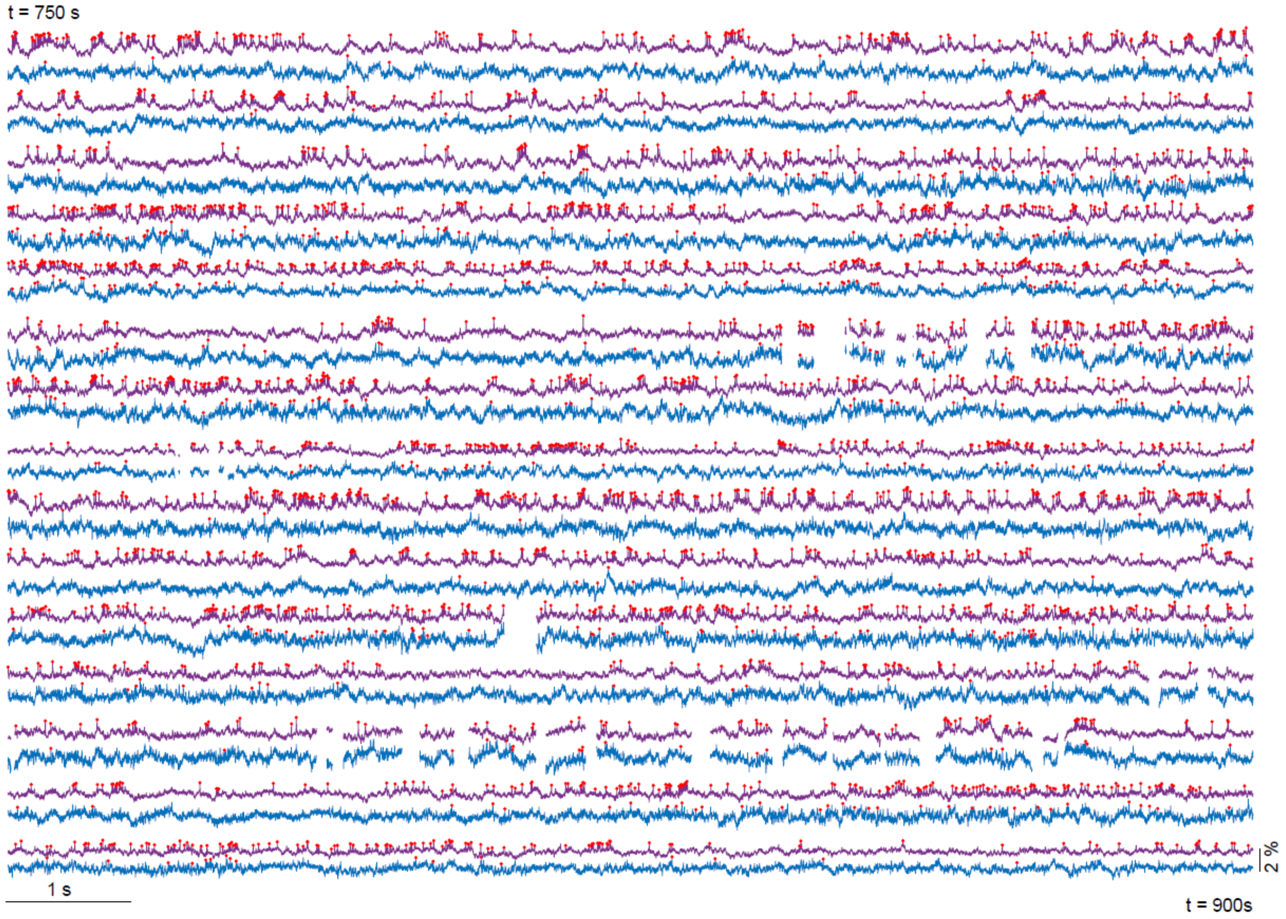


**Supplementary Figure 1**: An example 15-minute recording session with 2 simultaneously recorded neurons. Vm traces of the neurons are shown in blue and purple respectively. Red asterisks denote identified spikes. Time points with significant image motion were excluded.

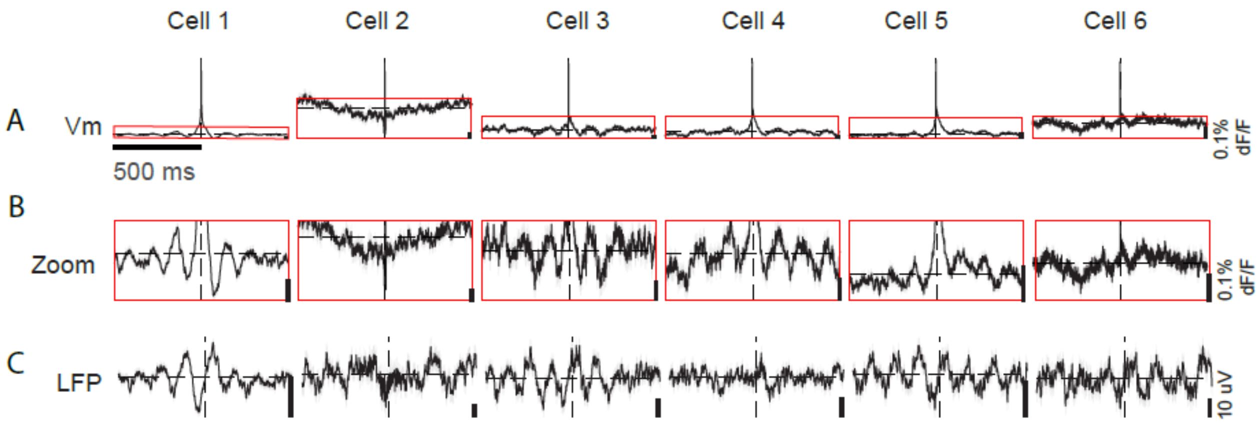


**Supplementary Figure 2**: **Spiking predominantly occurred on the rising phase of Vm and LFP theta oscillations**. An example session showing (A) Vm of a neuron aligned to the identified spikes. (B) Zoom-ins of A. (C) LFP aligned to the identified spikes for each neuron. Black lines are mean, and shaded areas are SEM. Dashed lines indicate 0.

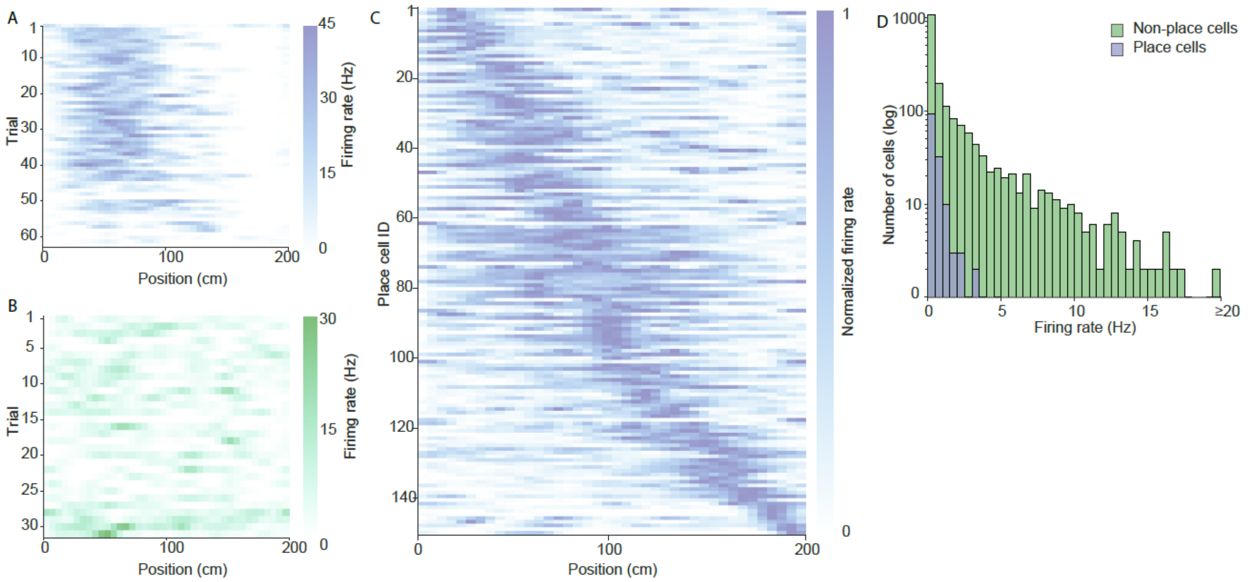


**Supplementary Figure 3: Place cells showing spatially specific spiking.** Example rate map of (A) a place cell and (B) a non-place cell. (C) The identified place cells tiled the entire virtual track. Neurons were sorted by the location of their place fields. Only the neurons containing a single place field are included (n = 7 mice, 143 neurons). (D) Firing rate distribution of place cells and non-place cells.

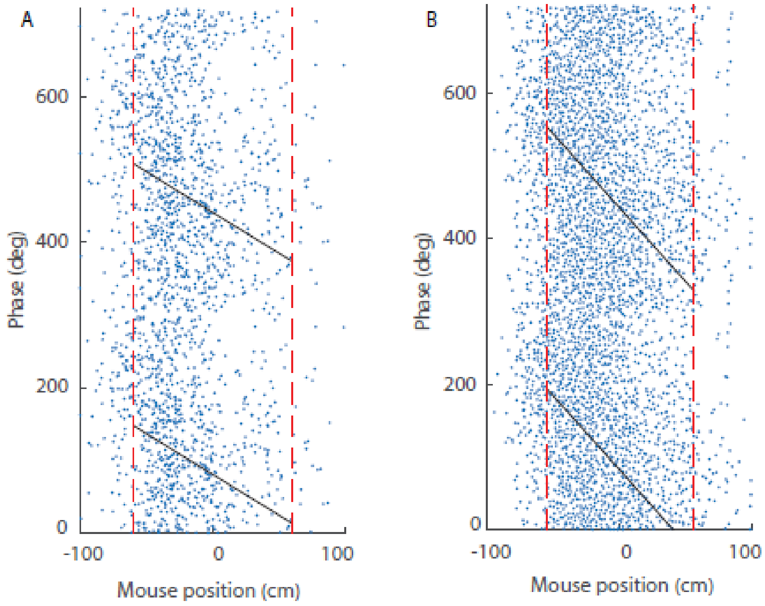


**Supplementary Figure 4: Example phase precession in two simultaneously recorded place cells.** (A) Spike timing relative to LFP theta phase. Black lines are the best fit linear-circular regression. The vertical red dashed lines mark the beginning and end of the place field. X-axis is centered to the place field. Y-axis shows two theta cycles. B) Same as in A, for a different neuron.
.

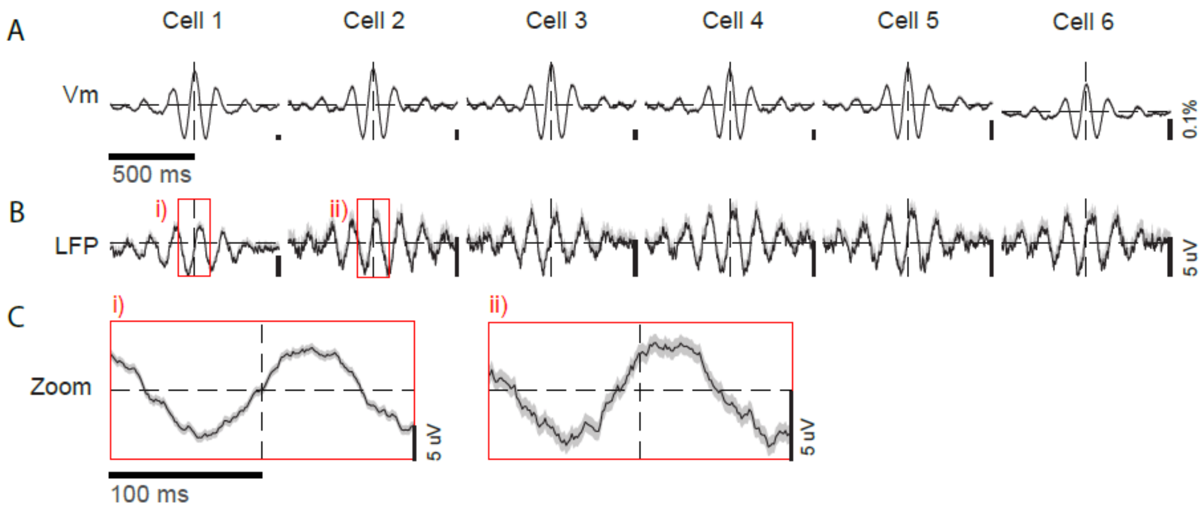


**Supplementary Figure 5**: **Vm theta peak in many cells occurred at specific phases of LFP theta**. (A) An example session showing Vm of simultaneously recorded neurons aligned to the peak of their own Vm theta. (B) LFP aligned to the identified spikes of each neuron. (C) Zoom-ins of B. Black lines are mean, and shaded areas are SEM. Dashed lines indicate 0.

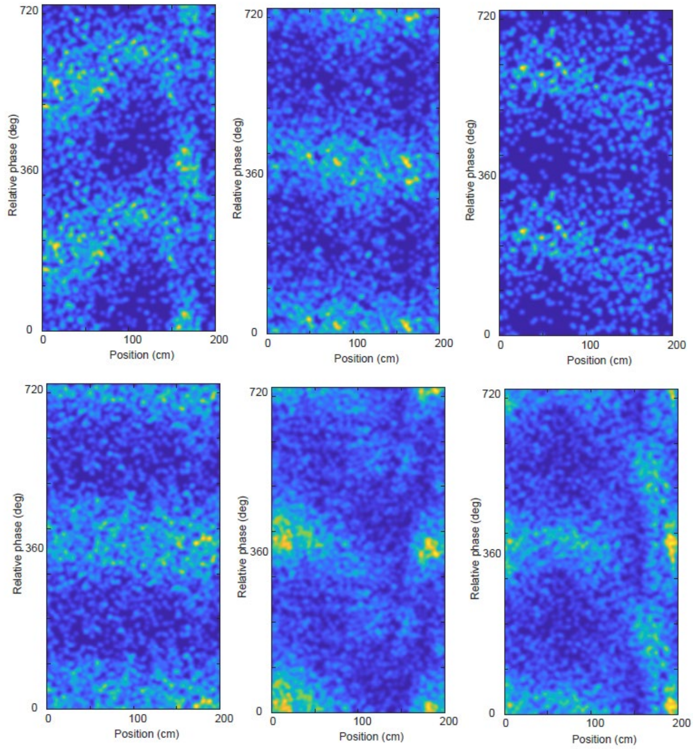


**Supplementary Figure 6**: Additional examples of the **relative phase of neuron pairs in Cluster A that dynamically changed along with spatial position**. Six additional example 2D histogram heatmaps for neuron pairs exhibiting gradual shifting of their relative phase over spatial position. These are additional examples from the same cluster as the six cell pairs shown in Figure 4A.

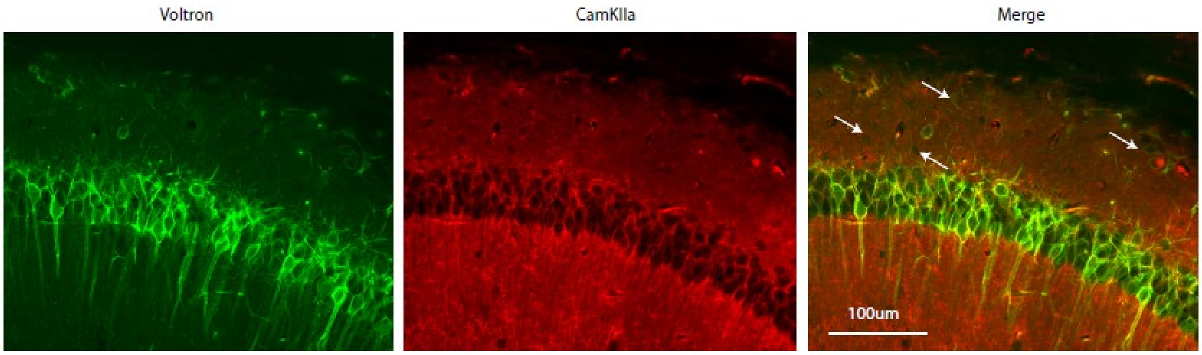


**Supplementary Figure 7**: **Voltron2 non-specifically labeled CA1 neurons**. Left, a representative confocal image showing Voltron 2 expression (Green). Middle, immunofluorescence of CamKIIa (red). Right, merge. Arrows indicate cells in the Stratum Oriens, that are positive or voltron2 but not for CamKIIa immunofluorescence.
